# Delineating In-Vivo T1-Weighted Intensity Profiles Within the Human Insula Cortex Using 7-Tesla MRI

**DOI:** 10.1101/2024.08.05.605123

**Authors:** C. Dalby, A. Dibble, J. Carvalheiro, F. Queirazza, M. Sevegnani, M. Harvey, M. Svanera, A. Fracasso

**Author notes:** **Corresponding author** Alessio Fracasso Hillhead Street 62KTDKT G12 8QE University of Glasgow, Scotland UK.

## Abstract

The integral role of the insula cortex in sensory and cognitive function has been well documented in humans, and fine anatomical details characterising the insula have been extensively investigated ex-vivo in both human and non-human primates. However, in-vivo studies of insula anatomy in humans (in general), and within-insula parcellation (in particular) have been limited. The current study leverages 7 Tesla magnetic resonance imaging to delineate cortical depth intensity profiles within the human cortex. Our analysis revealed two separate clusters of relatively high and low signal intensity across the insula cortex located in three distinct compartments within the posterior, anterior-inferior, and middle insula. The posterior and anterior-inferior compartments are characterised by elevated T1-weighted signal intensities, contrasting with lower intensity observed in the middle insular compartment, compatible with ex-vivo studies. Importantly, the detection of the high T1-weighted anterior cluster is determined by the choice of brain atlas employed to define the insular ROI. We obtain reliable in-vivo within-insula parcellation at the individual and group levels, across two separate cohorts acquired in two separate sites (n1 = 21, Glasgow, UK; n2 = 101, Amsterdam, NL). Results are further confirmed by deriving cortical depth dependent profiles from T1Map and R1Map images. These results reflect new insights into the insula anatomical structure, in-vivo, while highlighting the use of 7 Tesla in neuroimaging with potential implications for individualised medicine approaches.

**Graphical Abstract:** 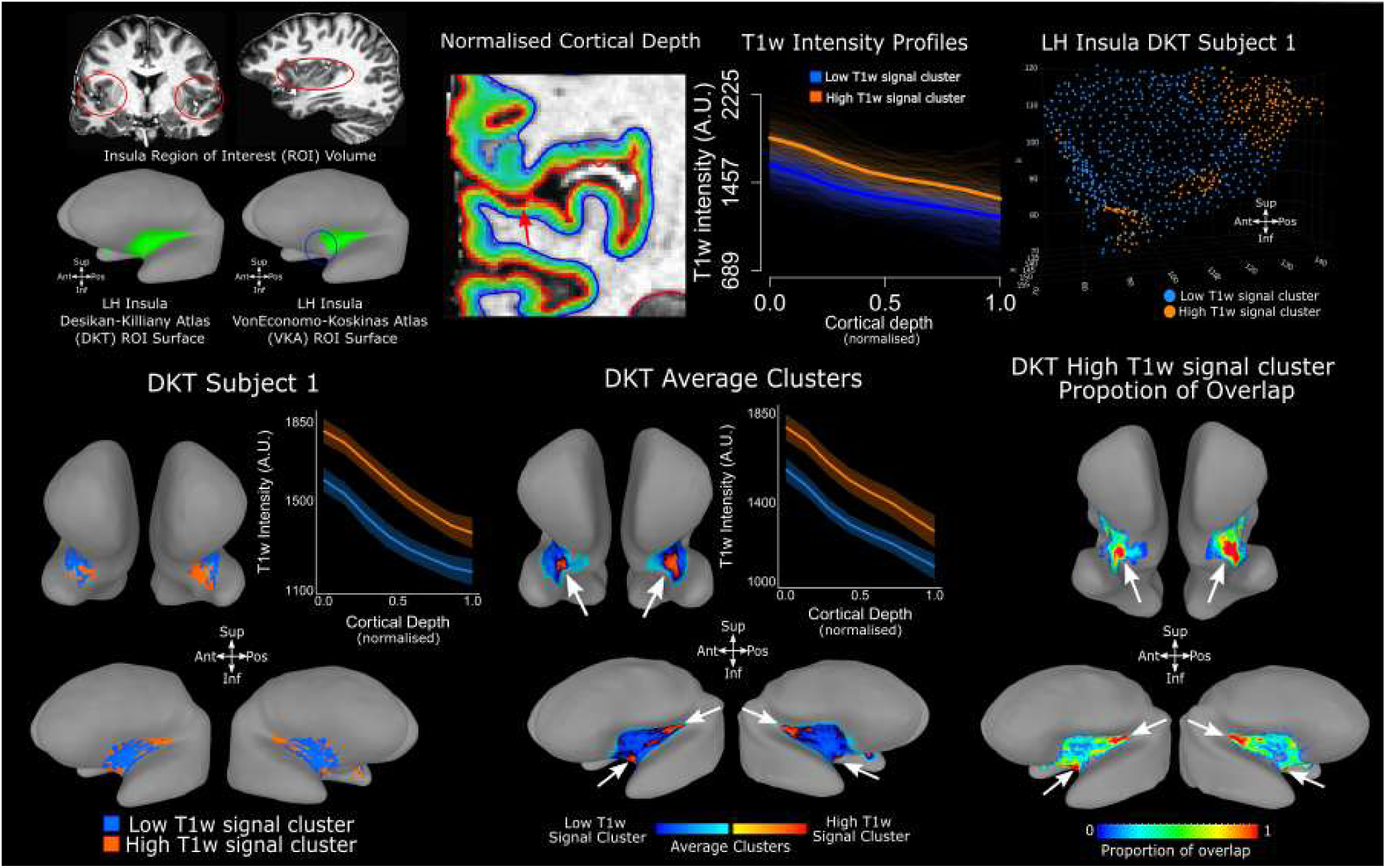

## Introduction

Folded deep within the lateral sulcus of each hemisphere, the insula cortex stands as a cornerstone of human function and behaviour (Menon et al., 2020; Gasquoine, 2014). It orchestrates activity that spans the domains of sensory perception (interoception, pain, gustation), emotional regulation (empathy, social emotions), and cognitive functions (goal-directed tasks, risk, memory and language) (Craig, 2002; Craig, 2004; Critchley et al., 2004; Critchley & Garfinkel, 2017; Critchley & Harrison, 2013). Such diversity in function has even prompted speculation about the insula’s candidacy for a central role in consciousness and self-awareness (de Haan et al., 2021; Medford & Critchley, 2010; Tisserand et al., 2023).

Anomalies in the insula’s function have been linked with clinical disorders such as schizophrenia, frontotemporal dementia, addiction, chronic pain and inflammation (Segerdahl et al., 2015; Droutman et al., 2015; Gebhardt & Nasrallah, 2023). Advancing our comprehension of the insula structure promises insights into both the normative landscape of human behaviour and the mechanistic, diagnostic, and therapeutic dimensions of clinical populations (Benarroch, 2019).

Over the years, several attempts have been made to parcellate the insula into smaller subdivisions. Early attempts relied on Brodmann’s seminal work on the distribution of neuronal bodies within human grey matter (cytoarchitectonics) (Brodmann, 1909; Geyer & Turner, 2013; Kurth et al., 2010a, Quabs et al., 2022). Subsequent research draws parallels with macaque cytoarchitecture, categorising the insula into granular, dysgranular, and agranular zones for posterior, intermediate, and anterior regions, respectively (Evrard, 2019; Nieuwenhuys, 2012a; Uddin et al., 2017). Similarly, immunohistochemical staining on macaque tissue by Gallay et al. (2012) revealed a ventral-to-dorsal increase in myelination. More recently, structural and functional distinctions between the insula’s posterior, ventral anterior, and dorsal anterior regions have been identified - the tripartite insula subdivisions (Menon et al., 2020; Uddin et al., 2017). Utilising diffusion MRI and functional connectivity, researchers have identified white matter tracts that connect each component of the tripartite insula subdivisions to frontal, limbic, and sensorimotor regions, facilitating the differential sensory, emotional, and cognitive functions, respectively (Klugah-Brown et al., 2023; Kurth et al., 2010b; Menon et al., 2020; Nomi et al., 2018; Uddin et al., 2017).

Furthermore, recent studies investigating the microstructure of the insula highlight the dynamic organisation of the region wherein information flows in a structured manner across its tripartite subdivisions. This process transitions from sensory-specific locations to transmodal locations that integrate diverse sensory, emotional and cognitive inputs moving along the anterior-ventral direction (Evrard, 2019; Royer et al., 2020). Moreover, the observed overlap in activation and the crosstalk among intra-insula regions, as discussed by Evrard (2019) and Uddin et al. (2014), suggest that these interactions underscore the insula’s role as a central hub for the integration of sensory and emotional inputs.

Recent advancements, exemplified by Royer et al. (2020), have expanded on this work by employing human T1/T2-weighted image intensity (T1/T2-w) as a proxy for cortical myeloarchitecture – the study of myelinated fibres distribution within grey matter (Geyer & Turner, 2013; Stüber et al., 2014; Vogt, 1903). This novel approach unveiled two major gradients in myelination: one from posterior to anterior portions of the human insula (Royer et al., 2020), also linked to post-mortem human cytoarchitecture (Kurth et al., 2010a), and possibly mirroring a shift from sensory to affective functions (Uddin et al., 2014). A second myeloarchitectonic gradient was observed running in the dorsal-to-ventral direction within the human insula, potentially linked to shifts between attention and cognition (Molnar-Szakacs & Uddin, 2022; Royer et al., 2020).

In this paper, we contribute further evidence to the structural tripartite subdivision of the human insula by leveraging the higher signal-to-noise ratio and acquisition resolution that can be achieved with human high-field imaging (7T). We report results using sub-millimetre T1-weighted (T1-w) magnetisation prepared rapid acquisition gradient recalled echo (MP2RAGE) images as well as T1Map and R1Map contrasts as a proxy of myelination and iron (Fukunaga et al., 2010; Stüber et al., 2014). We implemented an analysis pipeline that allowed us to gain access to cortical-depth dependent information from the human insula (Dumoulin et al., 2018; Fracasso et al., 2016a, 2016b, 2018, 2021; Waehnert et al., 2014, 2016). We then derived cortical depth dependent profiles and used them to parcellate the human insula into separate clusters. We demonstrate remarkably stable parcellations at the individual subject level, for the left and right insula. We applied the same pipeline for two independent datasets, across different age-groups, acquired from two different sites and scanner vendors (n1 = 21, Glasgow, UK; n2 = 101, Amsterdam, NL).

## Methods

### Subjects

This study involved two cohorts. The first comprises 21 subjects (Glasgow: age range: 23-38, 10 male) recruited from Glasgow University pool of subjects. The second cohort includes 101 subjects from the AHEAD dataset, Amsterdam (age range: 18-80, 45 male, see Alkemade et al., (2020) for more details). Glasgow: all experimental procedures were approved by the local ethics committee at the School of Medical, Veterinary and Life Sciences of the University of Glasgow (reference number: 200180191 and GN19NE455).

Inclusion criteria specified healthy adults from any range with no underlying neurological conditions. All subjects provided informed consent. See Alkemade et al., 2020 for experimental procedures and ethics approval of the AHEAD dataset.

### MRI Acquisition

#### Glasgow

MRI data was acquired on a 7T Siemens Magnetom Terra system (Siemens Healthcare, Erlangen, Germany) and a 32-channel head coil (Nova Medical Inc., Wilmington, MA, USA) at the Imaging Centre of Excellence (University of Glasgow, UK). We collected T1-weighted MP2RAGE anatomical scans for each subject (0.625 mm isotropic, FOV = 160×225×240 mm^3^, 256 sagittal slices, TR = 4680ms, TE = 2.09ms, TI_1_ = 840 ms, TI_2_ = 2370ms, flip angle_1_ = 5°, flip angle_2_ = 6°, bandwidth = 250Hz/px, acceleration factor = 3 in primary phase encoding direction). Total acquisition time was 12 minutes.

#### Amsterdam

MRI data was acquired on Philips Achieva 7 MRI scanner (Philips Healthcare, Best, The Netherlands) and a 32-channel head array coil (Nova Medical Inc., Wilmington, MA, USA) at the Spinoza centre for Neuroimaging (Amsterdam, the Netherlands). T1-weighted scans for each subject were collected using a modified MP2RAGE sequence (MP2RAGEME, Caan et al., 2019), 0.7mm isotropic, FOV=205×205×164 mm3, 205 sagittal slices, TR=6720ms, TE=3.00ms, T_I_=670ms, TI_2_ = 3855ms, flip angle_1_ = 7°, flip angle_2_ = 6°, bandwidth = 405Hz/px, acceleration factor SENSEPA = 2. Total acquisition time was 16.30 minutes.

#### T1-w, T1Map and R1Map images

T1-w signal per se is not quantitative, and the value can vary greatly depending on the scanner, making it difficult to compare between estimates obtained in different sites. We opted for using T1-w signal derived from MP2RAGE sequences as these were readily available from our site in Glasgow. However, for the Glasgow site data (21 participants), we did not have access to the original MP2RAGE acquisition files, including the two separate inversion times, which would have allowed us to derive quantitative T1 images. For this reason, we performed the analysis on T1-w, R1Map and T1Map data provided in the AHEAD dataset acquired in Amsterdam, representing a replication and an extension of the results obtained from the Glasgow dataset. See Results for further details.

#### Image Processing and Analysis

Data processing was conducted using Freesurfer (https://surfer.nmr.mgh.harvard.edu/), AFNI /SUMA (https://afni.nimh.nih.gov/pub/dist/doc/htmldoc/index.html), nighres, (https://nighres.readthedocs.io/en/latest/) and R (https://www.r-project.org/).

For each subject, we first skull stripped the MP2RAGE image (Figure 1A), applying the AFNI function `3dSkullStrip` to the second inversion image. The skull stripped anatomy was processed with the `recon-all` Freesurfer pipeline. Freesurfer output was converted into a SUMA, using the command @SUMA_Make_Spec_FS for visualisation and processing (Figure 1A).

**Figure 1.**
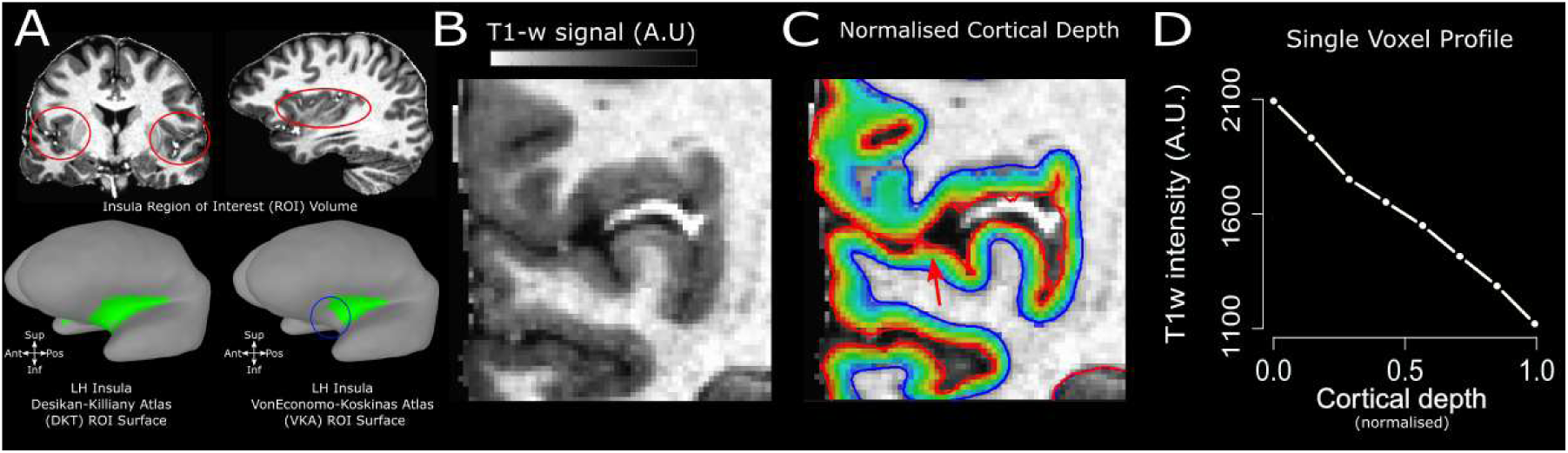
Pre-Processing. **A.** Insula ROI obtained from the DKT (no of voxels: 9587 mm^3^) and VKA (no of voxels: 5431 mm^3^) atlas. These atlases differ. Compared to the VKA, the DKT extends more towards the anterior part of the brain and dorsally with respect to the orbito-frontal cortex (see blue circle), and more towards the posterior part of the brain, towards the supramarginal gyrus. **B.** Coronal slice details of the MP2RAGE, single subject, left insula (0.7mm isotropic). **C.** Volumetric distance map along cortical depth. **D.** Individual T1-w profile passing through the voxel highlighted by the red arrow in panel D. T1-w signal is at its peak close to the white matter (0 on the x axis), then sharply decreases approaching the cerebro-spinal fluid (1 on the x axis).

#### Cortical Depth Analysis

We used nighres (Huntenburg, Steele and Bazin, 2018) to process the automatic segmentation of the anatomical (T1-w, MP2RAGE) images. Volumetric segmentations were obtained using the cruise algorithm (Han et al 2004), which yields a segmentation that is topologically correct and free from self-intersections corrected.

We generated white matter and grey matter level-sets using the nighres function `surface.probability_to_levelset()`. A volume-preserving distance map was computed between the white matter/grey matter boundary and the grey matter/cerebro-spinal fluid boundary in 8 separate level-set volumes (Figure 1C) using the nighres function `laminar.volumetric_layering()` using the equi-volume model. The equi-volume model provides a coordinate system of cortical depth which is independent from local cortical folding (Fracasso et al., 2018, 2021; Fabius et al., 2022; Waehnert et al., 2014, 2016).

To construct a cortical-depth profile for an individual voxel, we began at the levelset closest to the selected voxel (e.g. at the middle level set, corresponding to the middle cortical depth). We iteratively extended its surface normals outward toward the grey matter surface, storing intersection coordinates with each subsequent level set encountered. We then repeated this procedure in the opposite direction, projecting normals inward toward the white matter boundary. The two resulting coordinate streams were concatenated, forming a continuous cortical-depth profile associated with the selected voxel. Cortical depth dependent profiles were created by linearly interpolating signal intensity (T1-w, R1 and T1 map) along the profile’s depth coordinates (Fracasso et al., 2018; van Dijk et al., 2021a,b). Overall, each grey matter voxel was assigned to the profile of its nearest profile coordinate (see Figure 1D). We performed this procedure from all the voxels within grey matter, so a complete cortical depth-dependent profile, spanning between 0 and 1 along cortical depth, was assigned to each voxel within grey matter.

For this reason, an individual voxel at a cortical depth of say, 0.68, was associated with a complete cortical depth dependent profile, spanning between 0 (white matter) and 1 (grey matter surface) along cortical depth. This represented the cortical depth profile starting from white matter, passing through the individual voxel at cortical depth 0.68, and ending at the closest grey matter border.

Because adjacent voxels sample signals from nearly identical points along cortical depths, their profiles are highly similar and thus redundant. To reduce this redundancy, we retained only profiles originating from voxels with cortical depths between 0.15 and 0.85 for further analysis. Cortical thickness and cortical curvature were estimated using Freesurfer.

Please see supplementary material for further details on how we removed cortical curvature and cortical thickness contributions to individual cortical depth dependent profiles. (Supplementary Materials: *Removing Cortical Curvature and Cortical Thickness Contributions*).

#### Region Of Interest (ROI) Definition

We derived the insula ROI from two separate atlases to test the robustness of the proposed pipeline against different ROI definitions: i) the Desikan-Killiany atlas (Desikan et al., 2006), as it is widely used in the neuroimaging literature and ii) the Von Economo-Koskinas Atlas (Scholtens et al., 2018), as it has been used in a previous investigation focusing on characterising myelin gradients within the human insula (Royer et al., 2020).

#### Desikan-Killiany Atlas (DKT)

The atlas developed by Desikan et al., (2006) is a surface-based parcellation scheme for the human cerebral cortex. It was created using a dataset of 40 MRI scans to define cortical regions of interest (ROIs). The authors employed a semi-automated approach that combined manual labelling of specific cortical landmarks (sulcal representation) with automated parcellation algorithms (Fischl et al., 2004). The resulting atlas divides the cerebral cortex into 34 distinct ROIs per hemisphere, providing a standardised framework for structural and functional analyses. Later, the original Desikan-Killiany atlas was refined and a more detailed and consistent labelling protocol was developed, leveraging 101 brain images from the Mindboggle-101 dataset (Klein & Tourville, 2012). Regarding the insula cortex, the atlas defines a single insula ROI for each hemisphere without further subdivision into subregions.

#### Von Economo-Koskinas Atlas (VKA)

The Von Economo-Koskinas atlas, originally published in the early 20^th^ century, is a comprehensive brain atlas that parcellates the human cerebral cortex based on cytoarchitectonic differences observed in the cellular structure and organisation of the cortical layers (Economo & Koskinas, 1925). Created by Constantin von Economo and George Koskinas, this atlas was developed through meticulous microscopic examination of stained brain sections. The atlas divides the cerebral cortex into numerous cytoarchitectonic areas, each characterised by distinct patterns of cellular organisation and density. Regarding the insula cortex, the Von Economo-Koskinas atlas identified several cytoarchitectonic subdivisions within this region. The most notable are the agranular, granular and dysgranular insula. This granularity makes the atlas a valuable resource for understanding the structural organisation of the insula, which is critical for correlating anatomical features with function and clinical observations. Scholtens et al., (2018) made this historic atlas available by manually segmenting individual T1 scans and using FreeSurfer software to construct a digitised group-specific cortical parcellation atlas file. Royer et al.’s, (2020) findings were based on the updated version of this atlas (Scholtens et al., 2018), and we also employed the same version of the VKA atlas to facilitate comparisons with previous research (Figure 1A).

#### Differences between DKT and VKA Insula ROI

It became quickly apparent that the insula ROI defined from the DKT atlas differed from the insula ROI obtained from the VKA atlas. Namely, the former extends more towards the anterior part of the brain (Figure 1A), and dorsally with respect to the orbito-frontal cortex. Moreover, DKT insula ROI (9587 mm^3^) extends more towards the posterior part of the brain and the supramarginal gyrus compared with the VKA atlas insula ROI (5431 mm^3^).

This difference between the two atlases can be ascribed to the different criteria used to define ROIs in each atlas. The DKT atlas adopted a sulcal approach, based on manually tracing from one sulcus to another, incorporating the gyrus in between (Desikan et al., 2006; Klein & Tourville, 2012). On the other hand, the VKA atlas is based on the original work on

Von Economo and Koskinas (Economo & Koskinas, 1925; Scholtens et al., 2018), which relies on cytoarchitectonics: the distribution of neuronal bodies within human grey matter. Given the different criteria adopted in the definition of the two atlases, it is not surprising to notice differences in ROI boundaries. Here, we are interested in providing a characterisation of intra-insular parcellation using structural MRI along cortical depth. For this reason, we opted for reporting the results using both atlases (DKT and VKA), assessing potential differences between the two and allowing us to compare our results with the current literature (Royer at al., 2020).

#### Pre-processing cortical depth dependent profiles

For each subject and insula ROI (VKA and DKT), we loaded cortical depth dependent data data in R using the function `read.AFNI()` from the file AFNIio.R from AFNI. Our objective was to compare separate insular ROIs from different atlases. Each insula ROI for one participant encompasses approximately 20,000 voxels, each associated with an XYZ coordinate within the individual participant’s anatomical space. To streamline computations, we applied k-means clustering (k = 1,000) based on the voxel’s XYZ coordinates to uniformly subsample each insula ROI into clusters of roughly 10–20 neighbouring voxels. The use of kmeans clustering to parcellate human neocortex was pioneered by Geyer and colleagues (2011), applying it to the identification of early visual cortex. Importantly, no anatomical or other features guided the clustering, preserving an unbiased spatial representation. To increase signal-to-noise ratio and minimise the potential contribution of small grey matter segmentation errors, we divided each insula ROI in 1000 separate cortical location, each containing approximately 20 neighbouring voxels (range = 15-30). We averaged the cortical depth dependent profiles from each cortical location, thus obtaining 1000 averaged profiles for each subject and insula ROI. From now on, we will refer to the averaged profiles simply as profiles. Moreover, for each cortical location we obtained an estimate of cortical curvature and cortical thickness from Freesurfer.

#### Cluster Analysis

We ran a cluster analysis on the cortical depth dependent profiles (Figure 2). We adopted the k-means clustering implementation in R, using Euclidean distance as distance metric in cortical depth profile space in the *k-means* function. Please note that clustering was performed on cortical depth dependent profiles only, and the algorithm did not have access to the 3D coordinates -XYZ- of individual voxels. To determine the optimal number of clusters, we inspected the silhouette score of the clustering solution, using the function `fviz_nbclust`, from the library *factoextra* in R. For any given entry in the set, the silhouette score measures how similar an entry is to its own cluster compared to other clusters. The silhouette score ranges from −1 to 1. Values above 0 indicates that the entry is well matched to its own cluster and poorly matched to neighbouring clusters. The opposite is true for values below 0. The silhouette score for the entire dataset is the average of the silhouette scores of all individual entries. The plot of silhouette score across different number of clusters provides a way to visually assess the optimal number of clusters. We have generated a separate silhouette plot for each subject and insula (left and right hemisphere). Moreover, we derived the average silhouette plot from individual silhouette plots for a number of clusters ranging between 1 and 10.

**Figure 2.**
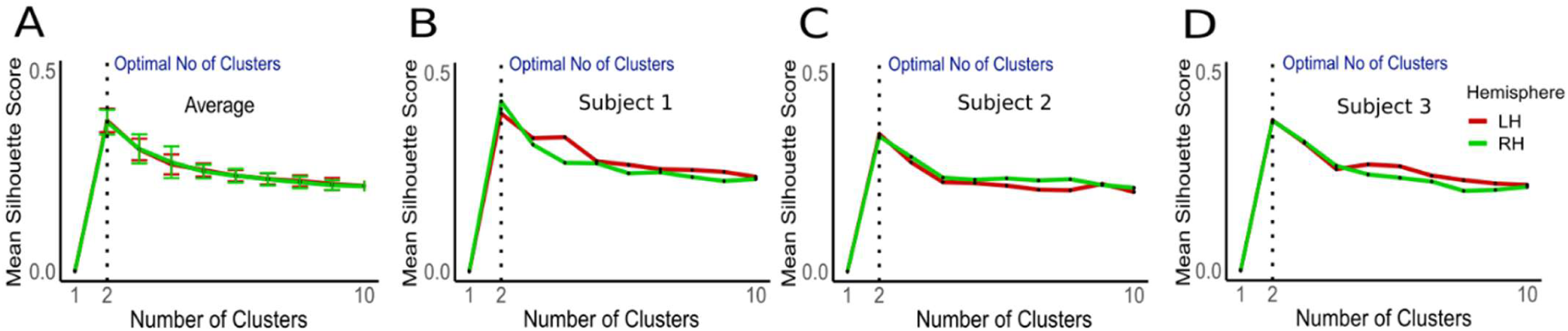
Silhouette Plots of K-Means Clustering Solution. **A.** Average silhouette plot for the left (red line) and right (green line) hemisphere DKT insula ROI across all 21 subjects, Glasgow dataset. Number of tested clusters ranged from 1-10. Dotted line indicates the optimal number of clusters (2). Error bars around each line indicate variability (+/- 1 standard deviation). **B-D.** Silhouette plots for three subjects to exemplify the optimal cluster number (2) remains consistent at the individual subject level.

#### Surfaces

We used the standard-mesh approach in SUMA (AFNI) to visualise data at the individual and average population level. A standard-mesh version of a surface is virtually an identical 3D mesh of the individual; however, this approach has the advantage that each node of the new mesh represents the same cortical location across subjects. The original subject’s mesh is recreated using the template mesh, instead of mapping a subject’s data value onto the template mesh (Saad & Reynolds, 2012). For each subject and insula ROI (VKA and DKT), we projected the k-means output sorted by the average intensity. We plotted the separate clusters on individual subjects’ surfaces using a categorical variable. Moreover, we plotted the proportion of overlap for each sorted cluster across the population. For each node in the insula ROI, we computed the proportion of subjects for which that specific node was labelled with the same cluster (ranked). This results in an overlap map, showing the consistency of insula parcellation across the population.

### Analysis

#### Stability of insula cluster maps

We assessed the stability of insula cluster maps using a leave-one-out procedure. For each iteration, we computed the average cluster map from n-1 individuals and tested the similarity between the n-1 average and the nth individual insula map using a logistic regression model and the logit function as link. For each individual and insula, we obtained the corresponding t statistic and p-value, representing how well the n-1 average cluster map captured the variability in the individual insula map. Building on the estimates of the logistic regression model, we computed the areas-under-the-curve (AUC) within a receiver-operating-characteristic curve (ROC) framework for each subject and insula. We report the AUC value and associated D-prime for each subject and insula in our dataset. D-prime is a measure derived from signal detection theory (Hautus et al., 2021) to quantify the ability to discriminate between two distributions, in this case between the distribution of separate clusters. D-prime represents the separation between the means of two distributions, standardised by their standard deviations. A value of 1 indicates a level of discriminability following approximately a hit-rate of 75% and a proportion of false-alarms of 30%.

#### Clusters or gradient

When discussing clusters in any modality, it is necessary to acknowledge that what might appear to be separate clusters, could instead be a continuous gradient. After thresholding by any clustering algorithm, a continuous gradient could be interpreted as separate clusters. On the other hand, ‘true’ separate clusters should be identified by a sudden change (a transient) in properties when transitioning from one cluster to another. See also the section: *Clusters or Gradients, simulations* in the Supplementary Materials.

To disentangle between the ‘cluster’ and the ‘gradient’ alternatives, we performed the following analysis on T1-w signal.

Inspecting clustering output at the T1-w profile level, it became apparent that different clusters were defined by different offsets. For this reason, for each subject and insula ROI, we projected on the MNI surface the T1-w profile intercept of each T1-w profile.

We reasoned that a gradient between the two separate clusters would manifest as a linear change in T1-w profile intercept along the underlying cortical distance between the clusters. On the other hand, a transient change in T1-w profile intercept along the cortical distance between clusters should be better approximated by a stepwise function. Please note that we use the term ‘cortical distance’ referring to the geodesic distance over the cortical surface, i.e. the shortest path between vertices over the cortical surface. We manually drew ROIs, crossing the border between neighbouring clusters on the MNI surface space and computed the corresponding geodesic distances within the ROI.

We computed the vertex-to-vertex geodesic distance from each node within the ROI – the adjacency matrix. We computed the spectral decomposition of the adjacency matrix and selected the first eigenvector of the decomposition. This vector represents a map of cortical distance over the ROI along its longest axis, where each point in the map is the geodesic distance of a node along the selected axis with respect to 0 – the middle of the ROI along its longest axis (Almeida et al., 2023; Fracasso et al., 2021; Grady & Polimeni, 2010; Lombaert et al., 2011). This cortical distance (or geodesic distance) provides a common reference frame against which we can test the linear (gradient) versus stepwise (cluster) hypothesis.

We plotted the T1-w profile intercept along the cortical distance and statistically assessed whether the data could be best fit by a linear or stepwise relationship. We tested a linear relationship fitting two parameters: an intercept and a slope (Eq. 1).

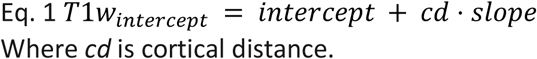

Where *cd* is cortical distance.

We used a cumulative Gaussian function with three parameters (multiplicative factor – mult -, shift and sigma, eq. 2) to test the stepwise alternative.

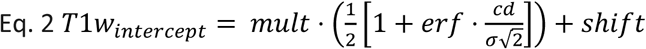

We compared the goodness of fit between the two models (linear and cumulative Gaussian) using the AIC criterion, penalising for the difference in the number of parameters between the two models (two against three). Statistical analyses and model fits were performed using R (https://www.r-project.org/).

#### AHEAD dataset

We applied the same processing and analyses described above to the T1-w images from the AHEAD dataset (Alkemade et al., 2020), as well as to the T1Map and R1Map images, which represent quantitative maps of T1, reflecting a proxy for myelin content (Bock et al., 2013; Glasser & Essen, 2011; Lutti et al., 2013).

## Results

### T1-w Cortical-Depth Dependent Profiles of the Human Insula

First, we present our results on the Glasgow dataset (21 subjects). These are followed by the results from the AHEAD dataset (Alkemade et al., 2020). For each dataset, here we present the results obtained from the DKT atlas, followed by the results of the VKA atlas. Following pre-processing, T1-w signal profile sampling (Figure 1A-D) and clustering (Figure 2), our analysis reveals two distinct profiles of low and high T1-w intensity, respectively (Figure 2). Qualitatively, using the DKT insula ROI, high T1-w intensity profiles tended to be localised in the anterior-inferior and posterior-superior insular locations (Figure 3).

**Figure 3.**
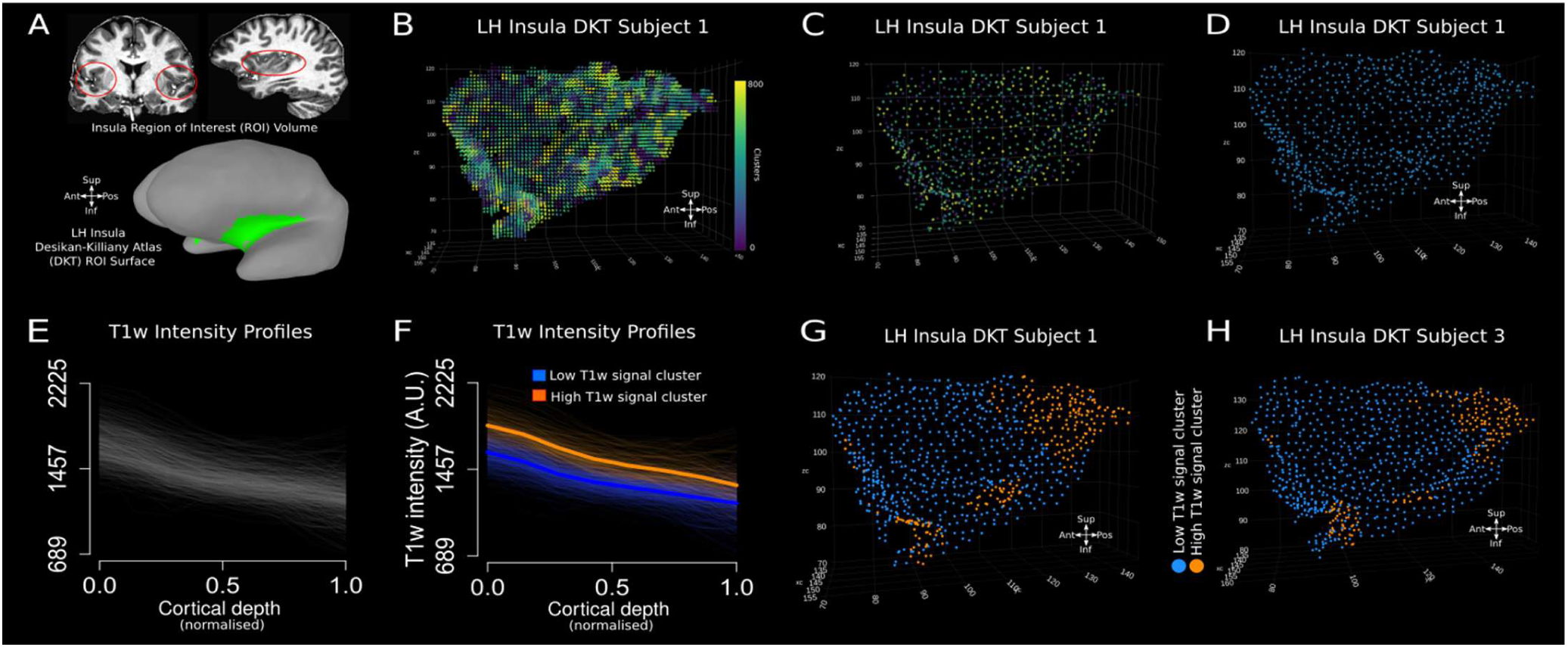
Individual Insula Clustering. **A.** Insula ROI obtained from the DKT atlas as viewed on the T1-w MRI volume (panel 1 and 2; red circles) and SUMA surface (green shading). **B**. Starting Insula ROI (approximately 20,000 voxels). Each voxel is assigned a different colour based on the right colour bar. **C.** Same insula ROI as in B, subsampled to 1000 points after clustering the XYZ coordinate of each voxel represented on the left (panel B). Each point represents the average of each coordinate within one of the 1000 clusters. **D.** Representation of the same insula as in panel C, using the same colour for each of the 1000 points. **E.** Individual T1-w profiles from the same insula from panel D. **F.** K-means clustering results based on the T1-w profiles intensity, separating high T1-w profiles from low T1-w profiles. Please note that a silhouette analysis indicated 2 as the optimal number of clusters. See the Results section: ‘Cluster Analysis on T1-w Profiles’. **G.H**. High T1-w profiles and low T1-w profiles colour coded on two individual subjects in native space (left insula). Qualitatively, in these two insula ROIs (from DKT atlas), the high T1-w intensity profiles tend to be localised in the anterior-inferior and posterior-superior insular locations.

### Cluster Analysis on T1-w Profiles

K-means clustering and silhouette plot (Figure 2) evidence the optimal number of clusters for dividing T1-w intensity profiles in the insula across both left and right hemispheres. For average (21 participants) and three individual subjects’ examples, the mean silhouette score peaks at two clusters and monotonically declines thereafter. This indicates that two clusters is the optimal solution at the individual insula level as well as across subjects. Should the optimal number of clusters be different (six clusters, for example), we would expect to see a peak in silhouette score at six, with a subsequent decrease, as the number of clusters increases.

In the following figure (Figure 3), we qualitatively report K-means clustering results, on two subjects.

### T1-w Intensity Profiles and Clusters on the DKT Insula ROI

Our analysis revealed distinct high- and low-T1-w intensity profiles across DTK insula ROI, at an individual (Figure 3 and Figure 4A, B) and group level (Figure 4C). T1-w signal monotonically decreases along cortical depth in both the high- and low-T1-w signal cluster (estimate = - 487.65, t=21.10, p<0.001). Average T1-w profiles along cortical depth for individual subjects shows how clusters differ predominantly on their offset (intercept with respect to the white matter border), rather than their slope (rate of change along cortical depth) (Figure 4A-C). Statistical analysis confirms this difference in intercept (estimate = 144.20, t=21.10, p<0.001) without a change in slope (estimate =-4.94 t=-0.43, p=0.67) between the high and low T1-w intensity profiles. Crucially, the profiles in the high T1-w signal cluster can be found in two separate cortical location along the DKT insula ROI: in the posterior-superior portion and the anterior-inferior portion of the DKT insula ROI (Figure 4C,E). On the other hand, the profiles in the low T1-w signal cluster can be found in the middle of the DKT insula ROI, between the high T1-w signal profiles in the posterior-superior portion and those in the anterior-inferior portion of the DKT insula ROI (Figure 4D). The location of high and low T1-w clusters is consistent between individuals. More than 70% of individuals show high T1-w intensity clusters localised towards the anterior-inferior and posterior-superior DKT insula ROI (Figure 4C,E white arrows) and the low T1-w intensity cluster is localised in the middle region of the DKT insula ROI (Figure 4D). Across the population, the high T1-w clusters location occupy 28% of the DKT surface insula ROI (here we defined a high T1-w cluster location across the population as those nodes on the cortical surface where more than 70% of individuals show a high T1-w intensity cluster).

**Figure 4.**
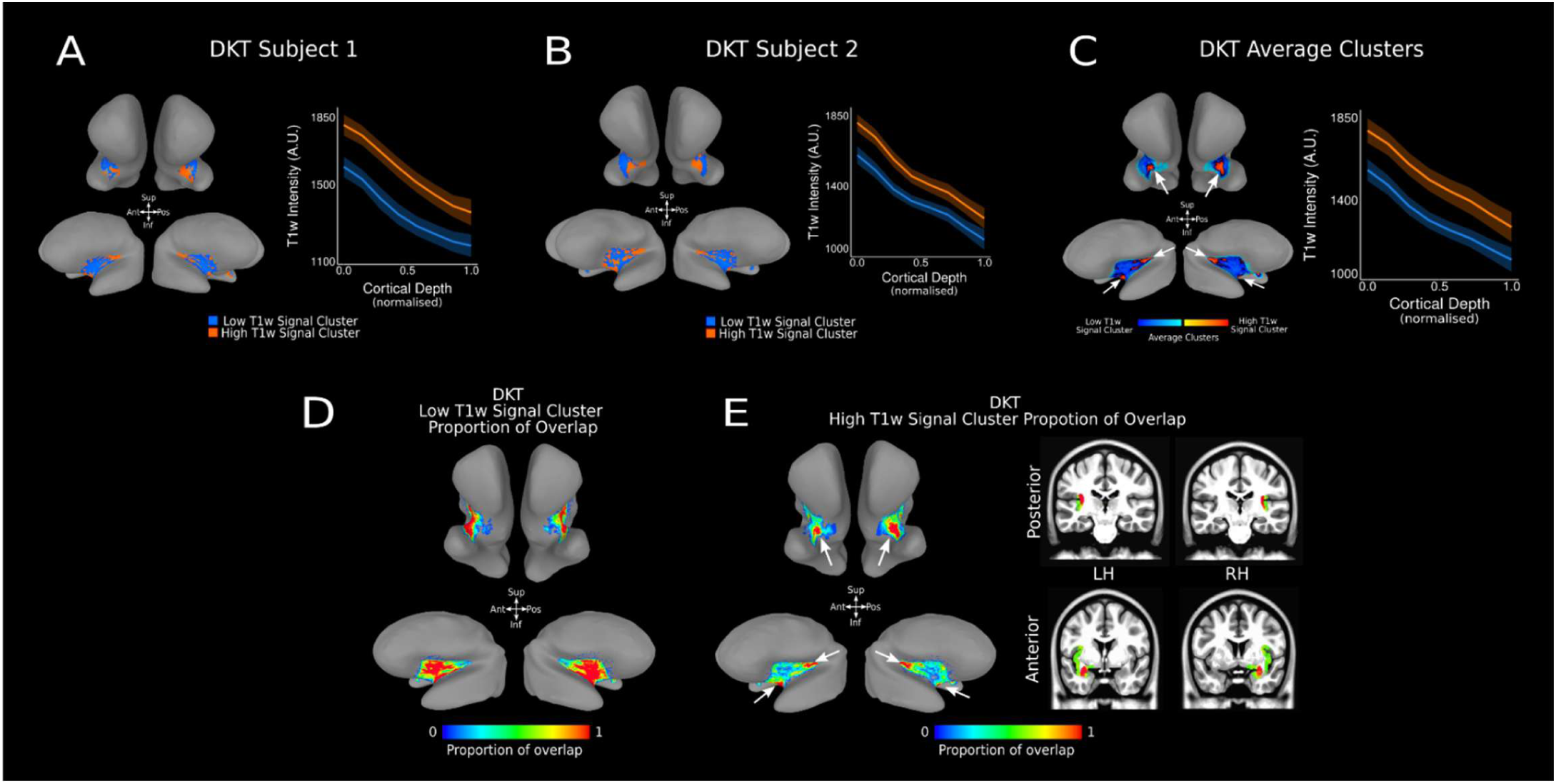
T1-w Intensity Profiles and Clusters in DTK insula ROI. A-B. Average high (orange) and low (blue) T1-w intensity profiles for the insula DKT ROI at the individual subject level. T1-w intensity is plotted along cortical depth and clusters are visualised on the MNI SUMA surface of the individual’s cortex (high T1-w intensity cluster – orange – and low T1-w intensity cluster – blue –). **C.** Average high (orange) and low (blue) T1-w intensity profiles for the insula DKT ROI across 21 subjects. T1-w intensity is plotted across cortical depth and average clusters across 21 subjects are visualised on the SUMA MNI surface. White arrows identify two areas of high T1-w signal clusters, located in the posterior-superior and anterior-inferior region of the DKT insula ROI. **D.** Proportion of overlap for low-T1-w signal cluster within the insula DKT ROI across 21 subjects, visualised on the average MNI SUMA surface. **E.** Proportion of overlap for high T1-w signal cluster for the insula DKT ROI across 21 subjects visualised on the MNI SUMA surface and on coronal slices in MNI space. White arrows identify two areas of high overlap for high-T1-w signal cluster, located in the posterior-superior and anterior-inferior region of the DKT insula ROI.

### T1-w Intensity Profiles and Clusters on the VKA Insula ROI

Following the results using the DKT atlas, we then aimed to replicate our findings in the VKA atlas employed by (Royer et al., 2020). Distinct high- and low-T1-w intensity profiles are also evident across the VKA atlas insula ROI at an individual level (Figure 5A,B) and group level (Figure 5C). As with the DKT atlas, T1-w signal monotonically decreases along cortical depth in both the high- and low-T1-w signal cluster (estimate = −469.40, t=-529.13, P<0.001). Equally, a high-T1-w intensity cluster is localised to the posterior-superior section of the VKA insula ROI (Figure 5C,E white arrows), while the low-T1-w intensity cluster is localised in the middle-anterior region of the VKA insula ROI. Statistical analysis confirms this difference in intercept (estimate = 209.66, t=268.50, p<0.001) and in slope (estimate =-61.60 t=-47.63 p<0.001) between the high and low T1-w intensity profiles.

**Figure 5.**
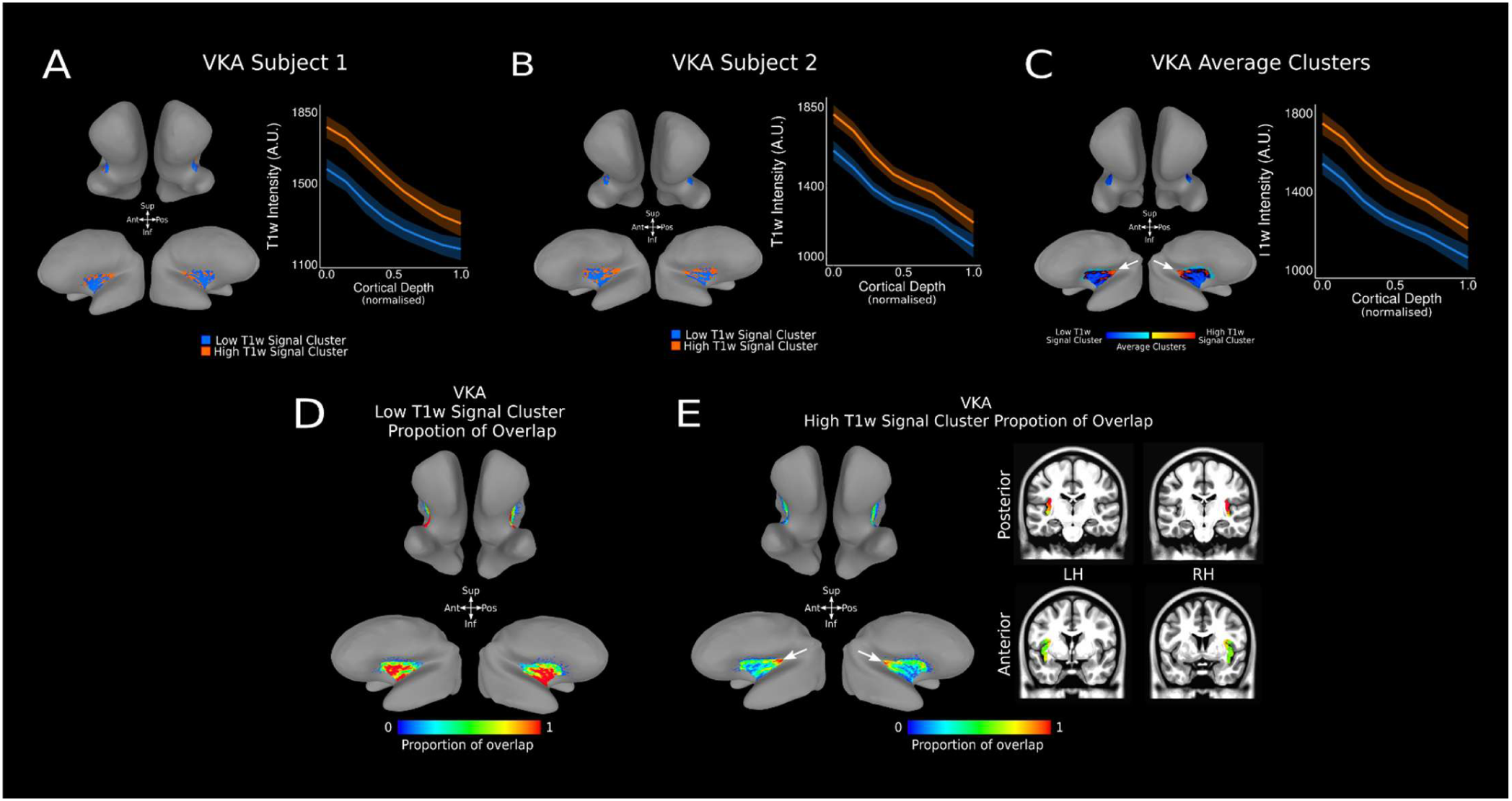
T1-w Intensity Profiles and Clusters in VKA Insula ROI. A-B. Average high (orange) and low (blue) T1-w intensity profiles for the insula VKA ROI at the individual subject level. T1-w intensity is plotted along cortical depth and clusters are visualised on the MNI SUMA surface of the individual’s cortex (high T1-w intensity cluster – orange – and low T1-w intensity cluster – blue –). **C.** Average high (orange) and low (blue) T1-w intensity profiles for the insula VKA ROI across 21 subjects. T1-w intensity is plotted across cortical depth and average clusters across 21 subjects are visualised on the SUMA MNI surface. The white arrow identifies the single area of high T1-w signal clusters, located in the posterior-superior region of the VKA insula ROI. **D.** Proportion of overlap for low-T1-w signal cluster within the insula VKA ROI across 21 subjects, visualised on the average MNI SUMA surface. **E.** Proportion of overlap for high T1-w signal cluster for the insula VKA ROI across 21 subjects visualised on the MNI SUMA surface and on coronal slices in MNI space. The white arrow identifies the single area of high overlap for high-T1-w signal cluster, located in the posterior-superior region of the VKA insula ROI.

### T1-w Intensity Profiles are Arranged in Clusters

It is important to note that any continuous gradient could result in separate clusters after being thresholded by any clustering algorithm. ‘True’ separate clusters should be identified by a sudden change (transient) in properties when transitioning from one cluster to another.

We manually drew ROIs, crossing the border between neighbouring clusters on the MNI surface space and computed the corresponding geodesic distances within the ROI (Figure 6 A,B). Different clusters are defined by different offsets (different T1-w profile intercept for the high- and low-T1-w signal clusters). For this reason, for each subject and cortical location within the insula ROI, we projected on the MNI surface the T1-w profile intercept (Figure 6C). A continuous gradient between the high-T1-w signal cluster and the low-T1-w signal cluster would manifest as a linear change in T1-w profile intercept along the cortical distance between the two clusters. On the other hand, a transient change in T1-w profile intercept along the cortical distance between clusters should be better approximated by a stepwise function (Figure 6D, blue and red curves for the linear and stepwise cumulative Gaussian fit, respectively).

**Figure 6:**
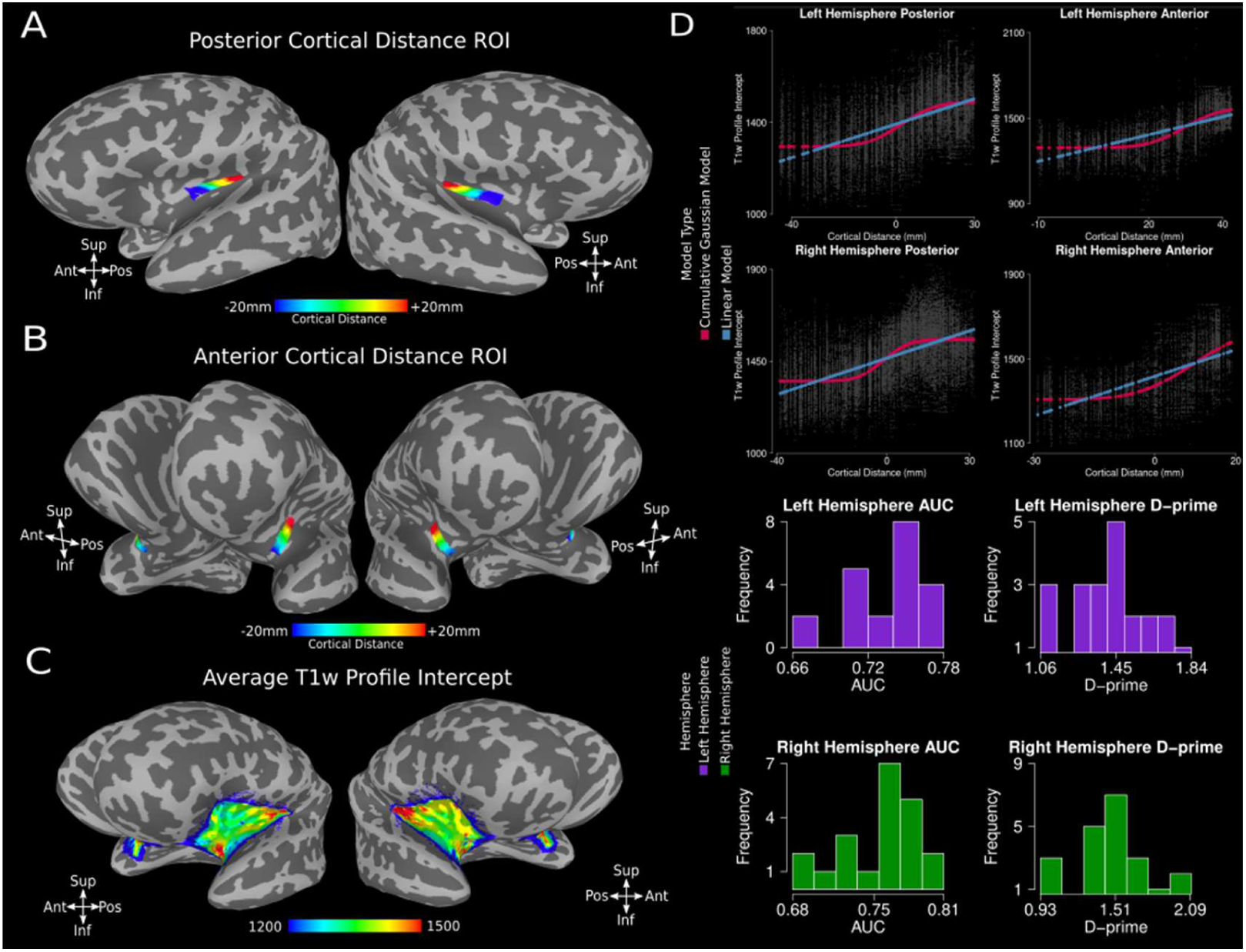
Gradient vs Clustering in DKT Insula ROI. **A.** Cortical distance maps for posterior insula (left and right hemisphere). **B.** Cortical distance map for anterior insula (left and right hemisphere). **C.** T1-w profiles intercept map over MNI SUMA surface for the left and right hemisphere. **D.** T1-w signal intensity intercept along cortical distance for each of the four ROI depicted in panels A,B. Original data is reported in grey, the result of the linear fit is reported in blue, the result of the cumulative Gaussian fit is reported in red (see legend). Histograms show the computed AUC and corresponding D-prime from the logistic regression analysis (see section Stability of insula cluster maps in Analysis). AUC values are above 0.5 in each individual participant, indicating the stability of the clustering results at the individual participant level. The corresponding D-prime values further support this observation, with the worst D-prime values ∼1 and the most representative value being ∼1.5, approximately corresponding to a hit-rate of 85% and a proportion of false-alarms of 25% in the leave-one-out scenario. See text for further details.

To disentangle between different alternatives, we fit a linear and a cumulative Gaussian model on T1-w signal intercept over cortical distance and assessed which model best captured variability in the data. Results indicate that the cumulative Gaussian model outperforms the linear model on each of the 4 ROI across high- and low-T1-w signal clusters in the posterior and anterior insula, for each hemisphere (see Table 1).

**Table 1.**
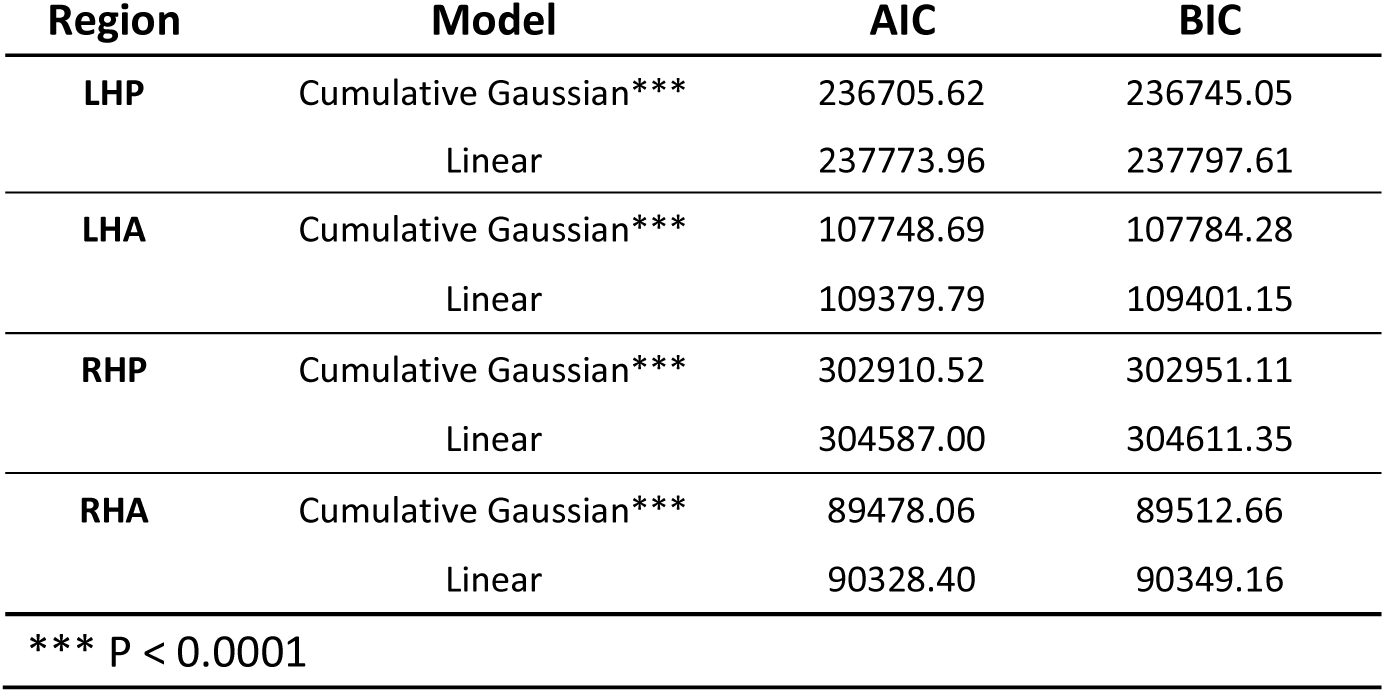
Summary Statistics of Linear and A Cumulative Gaussian Model on T1-w Signal Intercept Over Cortical Distance. Region refers to the regional location within the cortex (LH-: Left Hemisphere, RH-: Right Hemisphere, -A: Anterior, -P: Posterior). Model refers to the statistical model used to capture the variability for T1-w signal intercept over cortical distance. Akaike information criterion (AIC) and Bayes information criterion (BIC) values remain consistently lower for the Cumulative Gaussian model compared to the Linear Model across brain regions, indicating a better fit. Further statistical comparison using ANOVA supported these findings with significant results (see p-values) indicating the Cumulative Gaussian model more effectively captures the variability than the Linear model.

These results indicate that the clusters we report cannot be explained by a simple gradient along the insula, but rather by a transient change in properties along cortical distance (Figure 6D). A D-prime value of 1 indicates a level of discriminability following approximately a hit-rate of 75% and a proportion of false-alarms of 30%. This indicates that in the worst case, our leave-one-out model (the average of all individuals minus one) could correctly predict whether a high-T1-w profile of the left-out individual was categorised as being high-T1-w intensity 75% of the time, whereas a low-T1-w profile was wrongly categorised as high- on 30% of the times. We want to stress how our worst observed D-prime hovered around 1, with the most frequent value at the individual participant, individual insula level being ∼1.5, corresponding to a hit-rate of 85% and a proportion of false-alarms of 25%, indicating that these clusters can be separated reliably.

#### AHEAD Dataset

Utilising the same analysis pipeline, we replicated and expanded the reported initial findings from 21 participants (Glasgow) in a different cohort of 101 subjects using the AHEAD dataset. Importantly, the AHEAD dataset is a multi-contrast anatomical dataset where the authors acquired data using an MP2RAGE sequence with similar parameters as the MP2RAGE sequence we used to acquire data in Glasgow (see methods for more details about the dataset). Here we report results, from T1-w, T1Map and R1Map signal at the population level. Individual-level T1-w intensity profiles and clusters from the AHEAD dataset are reported in Supplementary Materials, Figure SF1.

### Group-Level T1-w Intensity Profiles and Clusters Within the DKT Insula ROI of the AHEAD Dataset

Utilising a larger sample from the AHEAD dataset resulted in a broader age range and subsequent increase in anatomical variability. To manage this variability in our group-level analysis, we divided the cohort into two age groups splitting the data in approximately two halves: ages 18-40 (n = 52 participants) and ages 41-80 (n = 49 participants). T1-w signal monotonically decreases along cortical depth in both the high- and low-T1-w signal cluster for participants aged 18-40 (estimate = −433.40, t=-710.20, p<0.001) and participants aged 41-80 (estimate = −446.13, t=-644.31, p<0.001). We observed distinct high- and low-T1-w intensity profiles across DTK insula ROI in both age groups (Figure 7A, B). Statistical analysis confirms a significant difference in intercept (I) and slope (S) in participants aged 18-40 (I: estimate = 249.87, t=398.68, p<0.001; S: estimate =-91.12, t=-88.19, p<0.001) and 41-80 (I: estimate = 227.73, t=363.49, p<0.001; S: estimate =-66.80, t=-64.62, p<0.001). The majority of individual subject’s low- (Figure 7C) and high- (Figure 7D) T1-w signal clusters are localised within the same cortical location of the DKT insula ROI. As in the Glasgow dataset, high T1-w intensity clusters are localised towards the anterior-inferior and posterior-superior DKT insula ROI (Figure 7B,D white arrows) and the low T1-w intensity cluster is localised in the middle region of the DKT insula ROI. Across the population, the high T1-w clusters location occupy 34% of the DKT surface insula ROI (here we defined a high T1-w cluster location across the population as those nodes on the cortical surface where more than 70% of individuals show a high T1-w intensity cluster).

**Figure 7.**
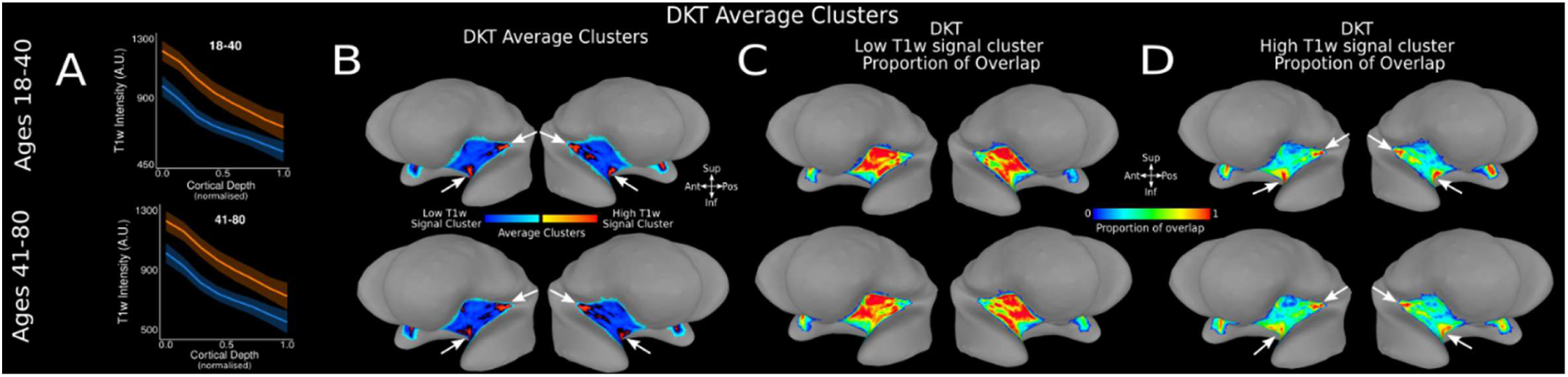
T1-w Intensity Profiles and Clusters in DTK Insula ROI for Two Age Groups in AHEAD Dataset. **A.** Average high (orange) and low (blue) T1-w intensity profiles plotted across cortical depth for the insula DKT ROI in age groups 18-40 (n=52) - panel 1 - and 41-80 (n=49) - panel 2. **B.** Average clusters across subjects in each age group are visualised on the SUMA MNI surface. White arrows identify two areas of high T1-w signal clusters, located in the posterior-superior and anterior-inferior region of the DKT insula ROI for each age group. **C.** Proportion of overlap for low-T1-w signal cluster within the insula DKT ROI across each age group, over on the average MNI SUMA surface. **D.** Proportion of overlap for high T1-w signal cluster for the insula DKT ROI across each age group visualised on the MNI SUMA surface. White arrows identify two areas of high overlap for high-T1-w signal cluster, located in the posterior-superior and anterior-inferior region of the DKT insula ROI.

### Group-Level T1-w Intensity Profiles and Clusters Within the VKA Insula ROI of the AHEAD Dataset

We replicate our findings employing the VKA atlas across the two AHEAD age groups. Distinct high- and low-T1-w intensity profiles are evident across the VKA atlas insula ROI at a group level for both age groups (Figure 8A). As with the DKT atlas, T1-w signal monotonically decreases along cortical depth in both the high- and low-T1-w signal cluster observed on the VKA insula ROI in participants aged 18-40 (estimate = −425.04. t=-1272.60, p<0.001) and participants aged 41-80 (estimate = −441.42, −1120.80, p<0.001. Similarly, statistical analysis confirm a significant difference in intercept (I) and slope (S) in participants aged 18-40 (I: estimate = 247.23, t=761.7, p<0.001; S: estimate =-114.61, t=-214.4, p<0.001) and 41-80 (I: estimate = 229.17, t=680.0, p<0.001; S: estimate =-76.46, t=-137.4, p<0.001). Moreover, a high-T1-w intensity cluster is localised to the posterior-superior section of the VKA insula ROI (Figure 8 B,D white arrow), while the low-T1-w intensity cluster is localised in the middle-anterior region of the VKA insula ROI.

**Figure 8:**
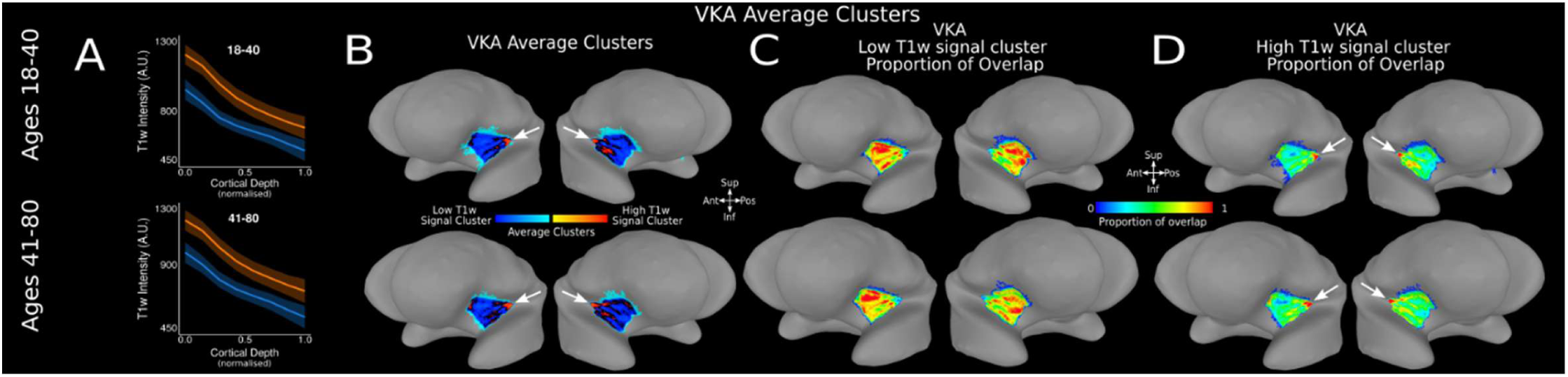
T1-w Intensity Profiles and Clusters in VKA Insula ROI for Two Age Groups in AHEAD Dataset. **A.** Average high (orange) and low (blue) T1-w intensity profiles plotted across cortical depth for the insula VKA ROI in age groups 18-40 (n=52) - panel 1 - and 41-80 (n=49) - panel 2. **B.** Average clusters across subjects in each age group are visualised on the SUMA MNI surface. White arrow identifies one area of high T1-w signal clusters, located in the posterior-superior of the VKA insula ROI for each age group. **C.** Proportion of overlap for low-T1-w signal cluster within the insula VKA ROI across each age group, visualised on the average MNI SUMA surface. **D.** Proportion of overlap for high T1-w signal cluster for the insula VKA ROI across each age group visualised on the MNI SUMA surface. White arrow identifies one area of high overlap for high-T1-w signal cluster, located in the posterior-superior of the VKA insula ROI.

### Group-Level T1Map and R1Map Intensity Profiles and Clusters Within the DKT Insula ROI of the AHEAD Dataset

We replicated our observations using T1Map and R1Map data available from the AHEAD dataset (Figure 9). T1Map and R1Map images, represent quantitative maps of T1, reflecting a proxy for myelin content (Bock et al., 2013; Glasser & Essen, 2011; Lutti et al., 2013). R1Map signal monotonically decreases along cortical depth in both the high- and low-signal clusters observed in the DKT insula ROI (estimate = −0.18. t=-860.4, p<0.001, Figure 9A). Note that higher R1Map values are indicative of a relatively higher myelination. Similarly, statistical analysis confirms a significant difference in intercept (I) and slope (S) (I: estimate = 0.14, t=550.20, p<0.001; S: estimate =-0.02, t=-45, p<0.001). T1Map results are qualitatively similar to those obtained from R1Map, although with inverted contrast, as T1Map signal monotonically increases along cortical depth (Figure 9B, t=460.5, p<0.001). For further reference, results from the average T1-w signal across the whole group (101 participants) are also reported in Figure 9.

**Figure 9:**
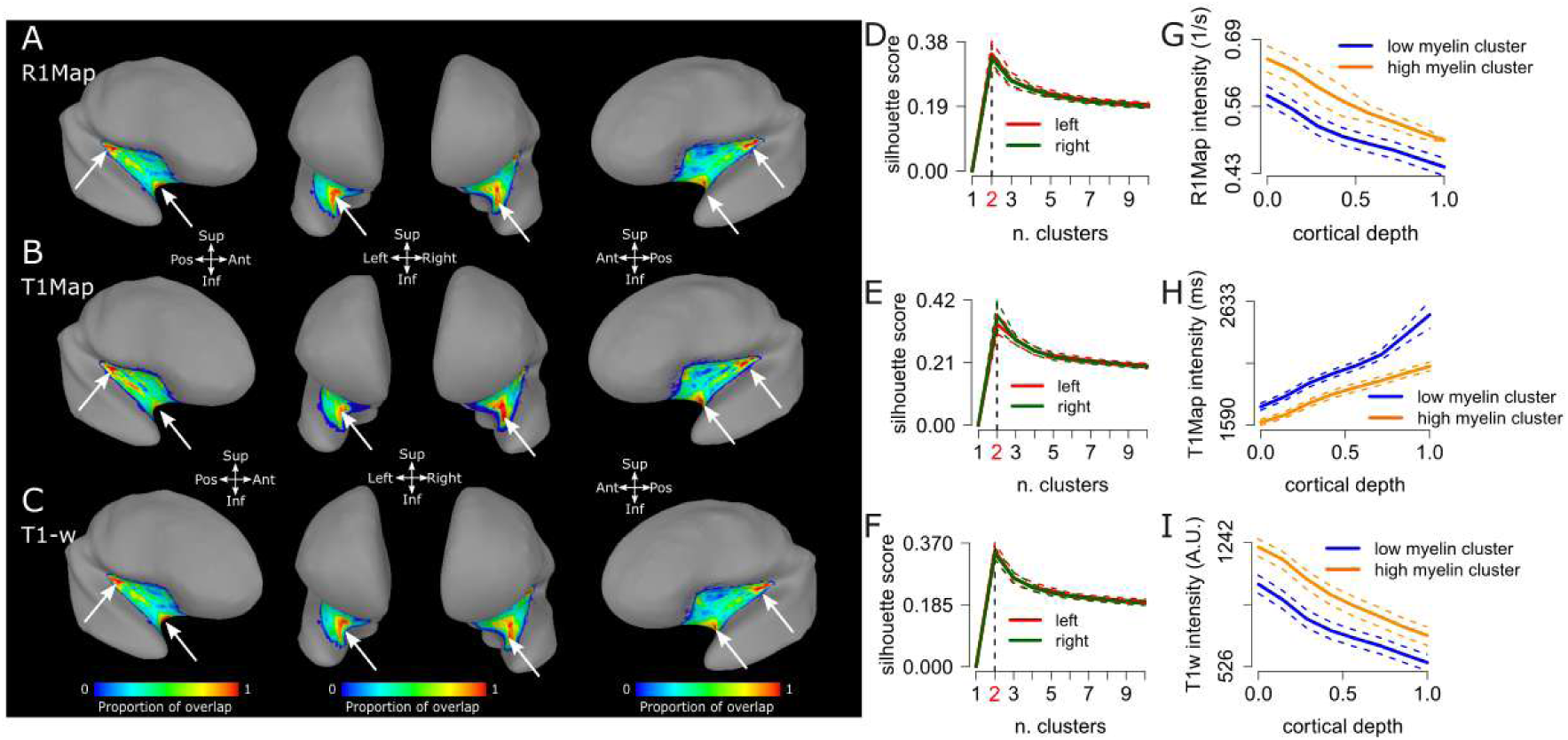
Clusters and Intensity profiles in DKT Insula ROI across different modalities (R1Map, T1Map and T1-w signal), AHEAD dataset, 101 participants. **A.** Proportion of overlap for high myelin cluster (relatively high R1Map signal) visualised on the MNI SUMA surface. Proportion of overlap for high R1Map signal cluster (corresponding to a relatively higher local myelination). White arrows identify areas of high overlap for high R1Map signal cluster, located in the posterior-superior and the anterior-inferior location of the DKT insula ROI. **B.** Same as in panel A, here reporting overlap maps for T1Map signal, where relatively low T1Map signal correspond to higher local myelination. **C.** Same as in panels A and B, here reporting overlap maps for T1-w signal, where relatively high T1-w signal corresponds to higher local myelination. **D-F.** Silhouette plots for the left and right hemisphere, derived from R1Map, T1Map and T1-w signal, indicating 2 as the optimal number of clusters. **G.** R1Map cortical depth dependent profiles clustered in relatively higher and lower myelination clusters, respectively. **H.** Same as G, here for T1Map signal. Note that in this case a lower T1Map signal correspond to a relatively higher myelination and vice-versa. **I.** Same as G and H, here for T1-w signal. Note that in this case a higher T1-w signal corresponds to a relatively higher myelination and vice-versa.

### Summary of Results

Our work offers novel insights into the parcellation of the insula. When employing the DKT atlas, our results reveal separate clusters in the anterior-inferior and posterior-superior insula locations. These findings are consistent across two cohorts, across different age-groups, acquired from two different sites and scanner vendors.

## Discussion

Cortical parcellation has been the target for neuroanatomists since the beginning of the twentieth century, to provide the first steps towards linking local cortical structure to the unknown underlying function (Economo & Koskinas, 1925; Geyer & Turner, 2013; Judas & Cepanec, 2010; Vogt, 1903). Initially, studies were performed ex vivo, on the basis of cortical differences in myeloarchitectonics (the study of myelinated fibres distribution within grey matter, along cortical depth (Judas & Cepanec, 2010; Vogt, 1903) and cytoarchitectonics (the study of neuronal bodies distribution within grey matter, along cortical depth (Brodmann, 1909). Today, we can take advantage of modern high field imaging and gain access to cortical structure with an unprecedented level of detail.

Here we applied techniques that allows us to gain insights on in-vivo parcellation, not only focusing on between-cortical area distinction, but showcasing an example of within-insula parcellation. Furthermore, we show how the within-insula parcellation is robust at the individual and group levels. We describe in-vivo cortical depth-dependent profiles of the insular cortex, drawing upon two independent datasets. We use two atlases to localise the insular cortex: the Desikan-Killiany Atlas (DKT) (Desikan et al., 2006) provided by Freesurfer, and the VonEconomo-Koskinas Atlas (VKA) (Scholtens et al., 2018), as employed in Royer et al (2020). The insula ROI from DKT atlas, extends more towards the anterior portion of the brain compared to the VKA atlas. We identify distinct clusters within the insula. These clusters are characterized by distinct offsets in T1-w, T1Map and R1Map signal at the level of the individual profile along cortical depth and showcase the possibility of performing within-area parcellation using high field imaging, in-vivo.

### Comparison with Previous Research

Previous studies on the insula focused on macaque and human cytoarchitecture reported by Evrard et al (2019) and Kurth et al (2010a). In macaques, Evrard et al. found distinct cytoarchitectonic features in both posterior and anterior insula. In the posterior dysgranular compartment, Evrard et al (2019) observed a series of thin, horizontally arranged stripes differing in cell density and laminar structure. On the other hand, in the anterior agranular compartment, the authors observed a localised cluster of large projection von Economo neurons. Kurth et al (2010a), report similar clusters of cytoarchitecture in the posterior region of the human insula but their analysis does not extend towards the anterior portion of the insula. Consistent with these observations, Royer et al’s (2020) seminal study used 3T HCP data (Glasser et al., 2013) to identify two primary myelin content gradients within the human insula at a group level: an anterior-to-posterior gradient and a dorsal-to-ventral gradient. This structural variation is also supported by a functional parcellation of the insula, linking differential myelin distribution to its diverse cognitive and emotional processing functions (Uddin et al., 2017).

In our current study, we used 7T MP2RAGE data employing the VKA atlas (as used in Royer et al., 2020) and observe similar increases in T1-w signal in the posterior-superior region of the insula. Moreover, employing the DKT atlas, we also detect an increase in T1-w signal in the anterior-ventral region, which was not previously reported in macaque studies. Two, non-mutually exclusive explanations could account for the discrepancy between our results and the known macaque insular organization. First, the anterior-inferior cluster may represent a genuine inter-species difference - present in humans but absent in macaques. Second, the divergence may reflect differences in anatomical definitions of the insula, akin to the discrepancy between the DKT and VKA atlases, the former of which extends the insular boundary further anteriorly. We believe our results indicate that the second alternative is more probable than the first. Specifically, the differences between atlases in the way the insula ROI is defined can lead to potentially missing the presence of the highly myelinated cluster in the anterior-inferior portion of the insula.

As illustrated by Figure 6, our findings evidence separate clusters characterized by relatively high myelination. On the other hand, Royer et al., (2020) instead documented the presence of a myelin gradient between comparable regions of the human insula. The difference between a continuous gradient reported in Royer et al., (2020) and the separate clusters reported here can be probably ascribed to the single subject analysis approach adopted in the current manuscript, as well as the isotropic, submillimetre spatial resolution of 7T MP2RAGE data. 7T MRI showcases several theoretical benefits over lower field strengths: increased spatial resolution, contrast- and signal-to-noise ratios (CNR and SNR, respectively). Overall, these benefits result in higher quality imaging within comparable acquisition times, which are manifest from a research and clinical perspective (Alkemade et al., 2020; Inglese et al., 2018; Isaacs et al., 2020; Kraff et al., 2015; Peerlings et al., 2019; Trattnig et al., 2018). Importantly, our approach allows for reliable parcellation at both the individual participant and group levels within the insula.

Previous research has delineated key structural characteristics of the posterior insula, including three distinct cytoarchitectural areas (Kurth et al., 2010a), high myelination (Royer et al., 2020), and extensive connections to other cortical regions such as the cingulate, parietal, frontal, and sensorimotor areas of the brain (Nomi et al., 2018; Uddin et al., 2017). These structural features underscore the posterior insula’s role in aggregating sensory information and facilitating functions such as temperature and pain perception (Nomi et al., 2018; Uddin et al., 2017). Our observed increase myelination in the posterior insula, as identified using both the DKT and VKA atlases, corroborates Royer et al.’s findings and provides additional structural insights into the insula structural organization. Tian & Zalesky (2018) identified meaningful gradients along the insula anterior posterior dimension based on the analysis of human resting state data. These results are compatible with the main anterior-posterior gradient reported in Royer et al., 2020 using structural MRI. Similar analytical approaches can be found in Farrugia et al., (2024) and Bajada et al., (2020). See also Supplementary materials: *Functional gradients in the human insula*.

Nomi et al. (2018) and Uddin et al. (2017) further elaborate on the connections between the anterior insula location and the frontal and limbic regions of the brain, emphasising the importance of these connections in supporting and influencing higher-level cognitive and affective processes. Based on the current literature, we speculate that the relatively high-myelination cluster we observe in both the posterior and anterior insular regions (using the DKT atlas ROI), could represent input and output zones within the insula’s network (Nomi et al., 2018; Uddin et al., 2017; Evrard et al., 2019). Through well-myelinated afferent and efferent connections, these regions might facilitate the flow of information to and from other regions within the insula and the surrounding cortical areas. This structural specialisation may be instrumental in supporting the diverse functionality attributed to the posterior and anterior insula regions, with the mid insula, as noted by Nomi et al. (2018) and Uddin et al. (2017), potentially acting as a transitional region. Furthermore, our findings also align with earlier reports of neuromodulatory and sensory input via connections to sensorimotor regions, thus underscoring its role as a hub for sensory integration (Gogolla, 2017; Klugah-Brown et al., 2023).

### Methodological Considerations/ Limitations

#### DKT and VKA Atlases

Our results indicate that the distribution of clusters is dependent upon the atlas used to define the insula. Starting from the DKT atlas, we identify two distinct clusters of relatively high- and low- myelination within the insula. Further analysis reveals how these clusters are localised in three distinct compartments within the insula ROI, distributed across its posterior, anterior-inferior, and middle sections, with the former two regions notably associated with relatively higher myelination levels.

We identify similar clusters starting from the VKA atlas. However, these are distributed in two cortical compartments instead, associated with high- and low-myelination clusters in the posterior and middle-anterior portion of the insula, respectively. A second high-myelination location cannot be identified in the anterior portion of the VKA insular ROI, due to the lesser extend of the VKA insula ROI towards the anterior portion of the brain compared to the DKT insula ROI.

This difference between the results obtained with the DKT and VKA atlases stems from the distinct criteria employed to delineate regions of interest (ROIs)—whether based on cytoarchitectonics or sulcal patterns—which inherently result in variations in ROI definitions across atlases.

Importantly, these differences can, in turn, lead to divergent characterizations of the functional or structural properties of a given ROI. Each atlas definition approach carries its own strengths and limitations. Given these atlas-specific idiosyncrasies, it is valuable to report ROI-based analyses using multiple atlases, as done in the current manuscript. This approach enables a more comprehensive understanding by highlighting both consistent findings and those that vary depending on atlas selection.

#### MRI contrast and its specificity for myelin

T1-weighted, T1Map and R1Map signals largely captures variances in lipid levels, a factor closely linked to myelin presence, yet the signal is also shaped by iron levels in the blood stream and iron-enriched lipids (Fukunaga et al., 2010; Koenig, 1991; Stüber et al., 2014). Consequently, the contrast observed in cortical depth dependent profiles may result from a mix of myelin and iron located within the grey matter.

#### Proximity to large arteries

As part of our analysis, we document that the anterior high myelination profile is adjacent to the nearby middle cerebral artery (MCA, Türe et al., 2000). See Supplementary Materials, Figure SF2 and section ‘*DKT Atlas, The Influence of The Middle Cerebral Artery on T1-w Signal Intensity*’. Given the spatial resolution of the MR image voxels to 0.7mm, our results are influenced by the partial volume effect whereby the presence of multiple tissues within a voxel leads to mixed signal intensities. This effect can cause inaccuracies in the representation of tissue boundaries and structures (Billot et al., 2020). Therefore, given the proximity, there is a possibility that the signal from the MCA could alter/confound the intensity of the anterior high myelination profile. As shown in our analysis (Supplementary Materials, Figure SF2), the average T1-w profile in the posterior insula cluster differs significantly from the average T1-w profile in the anterior insula cluster with the latter being shallower than the former. However, the proximity of the MCA does not disrupt the overall grey matter segmentation in neighbouring locations, nor does it disrupt the shape of the T1-w profile in the anterior insula cluster (DKT atlas, see Supplementary Materials, Figure SF2).

#### Clinical Relevance & Future Research Directions

Our results on the DKT atlas allow us to gain a deeper understanding of the structural features of the human insula and represent a structural counterpart to the functional tripartite insular parcellation (Menon et al., 2020).

Our observed two clusters solution (one cluster characterized by a relatively lower myelination compared to the second cluster) is arranged in 3 separate compartments over in the human insula: 1) the superior-posterior portion of the insula is characterized by relatively high T1-w intensity and high R1Map intensity, 2) the inferior-anterior portion of the insula is also characterized by relatively high T1-w intensity and high R1Map intensity, and 3) the middle portion of the human insula is characterized by a relatively low T1-w and low R1Map intensity.

Moreover, given their robustness at the individual level, they represent a promising first step towards an individualised – precision medicine - approach that could find venues of application for clinical populations where the insula appears to be critically involved.

For example, the insula’s activity correlates with the prolonged experience of pain, (Segerdahl et al., 2015).

We speculate that the study of individual myelination profiles could provide greater insights into the insula’s role in pain perception and modulation, possibly allowing to differentiate between healthy and pathological states, as well as enable more targeted diagnosis and treatment of patients with pain conditions. For example, as measured by PET, insula hypometabolism predicts remission with cognitive behaviour therapy and poor response to a selective serotonin reuptake inhibitor antidepressant (escitalopram), while insula hypermetabolism predicts remission with escitalopram and poor response to cognitive behaviour therapy (McGrath et al., 2013).

In this study, utilising high-resolution 7-Tesla imaging allowed us to step away from the traditional ’one-size-fits-all’ analytical approach, focusing instead on individual-specific data. This approach requires the development of novel software and analytical methods, tailored to manage this high-resolution data. By prioritising the unique characteristics of each subject’s brain structure, we ensure that the specific nuances are thoroughly captured, laying a foundation for personalised treatment planning (Waehnert et al., 2016).

## Conclusion

In conclusion, we describe in-vivo clusters in the anterior and posterior locations of the insula cortex on an individual and group level. We also characterise how application of different atlases can influence the identification of clusters within the insula. Building from historical approaches on cortical parcellation (Judas & Cepanec, 2010; Nieuwenhuys, 2012b; Vogt, 1903) and modern in-vivo imaging techniques (Alkemade et al., 2020; Bock et al., 2013; Glasser & Essen, 2011; Lutti et al., 2013; Royer et al., 2020), our work offers novel insights into the within-insula parcellation in-vivo, in humans. These findings offer a robust representation of in-vivo insula T1-w, R1Map and T1map signal used as a proxy of myelination, both at individual and group levels, and sets the stage for using insula cortical depth dependent profiles for clinical applications as chronic pain and inflammation (Segerdahl et al., 2015; Rolls, 2023).

## Statements and Declarations

Nothing to disclose.

## Acknowledgements

A.F. was supported by a grant from the Biotechnology and Biology Research Council (BBSRC, grant number: BB/S006605/1) and the Bial Foundation Grants Programme; Grant id: A-29315, number: 203/2020, grant edition: G-15516).

C.D was supported by a PhD grant by the Medical Research Council (MRC) as part of the Precision Medicine Doctoral Training Programme.

A.D was supported by a PhD grant from the Scottish Graduate School of Social Science, Doctoral Training Partnership (SGSSS-DTP), on behalf of the Economic and Social Research Council (ESRC, grant number: ES/P000681/1).

## Data availability

The Amsterdam Ultra-high field adult lifespan database (AHEAD) dataset can be found here: https://doi.org/10.21942/uva.10007840.v2

## References

Almeida, J., Kristensen, S., Tal, Z., & Fracasso, A. (2023). Contentopic mapping in ventral and dorsal association cortex: the topographical organization of manipulable object information. BioRxiv, 2023–11.

Alkemade, A., Mulder, M. J., Groot, J. M., Isaacs, B. R., van Berendonk, N., Lute, N., Isherwood, S. J., Bazin, P.-L., & Forstmann, B. U. (2020). The Amsterdam Ultra-high field adult lifespan database (AHEAD): A freely available multimodal 7 Tesla submillimeter magnetic resonance imaging database. NeuroImage, 221, 117200. 10.1016/j.neuroimage.2020.117200

Bajada, C. J., Campos, L. Q. C., Caspers, S., Muscat, R., Parker, G. J., Ralph, M. A. L., … & Trujillo-Barreto, N. J. (2020). A tutorial and tool for exploring feature similarity gradients with MRI data. NeuroImage, 221, 117140.

Benarroch, E. E. (2019). Insular cortex: functional complexity and clinical correlations. Neurology, 93(21), 932–938.

Billot, B., Robinson, E., Dalca, A. V., & Iglesias, J. E. (2020). Partial Volume Segmentation of Brain MRI Scans of Any Resolution and Contrast. In A. L. Martel, P. Abolmaesumi, D. Stoyanov, D. Mateus, M. A. Zuluaga, S. K. Zhou, D. Racoceanu, & L. Joskowicz (Eds.), Medical Image Computing and Computer Assisted Intervention – MICCAI 2020 (pp. 177–187). Springer International Publishing. 10.1007/978-3-030-59728-3_18

Bock, N. A., Hashim, E., Janik, R., Konyer, N. B., Weiss, M., Stanisz, G. J., Turner, R., & Geyer, S. (2013). Optimizing T1-weighted imaging of cortical myelin content at 3.0 T. NeuroImage, 65, 1–12. 10.1016/j.neuroimage.2012.09.051

Brodmann, K. (1909). Vergleichende Lokalisationslehre der Grosshirnrinde in ihren Prinzipien dargestellt auf Grund des Zellenbaues.

Craig, A. D. (2002). How do you feel? Interoception: the sense of the physiological condition of the body. Nature Reviews Neuroscience, 3(8), 655–666. 10.1038/nrn894

Craig, A. D. (2004). Human feelings: Why are some more aware than others? Trends in Cognitive Sciences, 8(6), 239–241. 10.1016/j.tics.2004.04.004

Critchley, H. D., & Garfinkel, S. N. (2017). Interoception and emotion. Current Opinion in Psychology, 17, 7–14. 10.1016/j.copsyc.2017.04.020

Critchley, H. D., & Harrison, N. A. (2013). Visceral Influences on Brain and Behavior. Neuron, 77(4), 624–638. 10.1016/j.neuron.2013.02.008

Critchley, H. D., Wiens, S., Rotshtein, P., Öhman, A., & Dolan, R. J. (2004). Neural systems supporting interoceptive awareness. Nature Neuroscience, 7(2), 189–195. 10.1038/nn1176

de Haan, E. H. F., Scholte, H. S., Pinto, Y., Foschi, N., Polonara, G., & Fabri, M. (2021). Singularity and consciousness: A neuropsychological contribution. Journal of Neuropsychology, 15(1), 1–19. 10.1111/jnp.12234

Desikan, R. S., Ségonne, F., Fischl, B., Quinn, B. T., Dickerson, B. C., Blacker, D., Buckner, R. L., Dale, A. M., Maguire, R. P., Hyman, B. T., Albert, M. S., & Killiany, R. J. (2006). An automated labeling system for subdividing the human cerebral cortex on MRI scans into gyral based regions of interest. NeuroImage, 31(3), 968–980. 10.1016/j.neuroimage.2006.01.021

Droutman, V., Read, S. J., & Bechara, A. (2015). Revisiting the role of the insula in addiction. Trends in Cognitive Sciences, 19(7), 414–420. 10.1016/j.tics.2015.05.005

Dumoulin, S. O., Fracasso, A., van der Zwaag, W., Siero, J. C. W., & Petridou, N. (2018). Ultra-high field MRI: Advancing systems neuroscience towards mesoscopic human brain function. NeuroImage, 168, 345–357. 10.1016/j.neuroimage.2017.01.028

Economo, C. F. von, & Koskinas, G. N. (1925). Die cytoarchitektonik der hirnrinde des erwachsenen menschen: J. Julius Springer Verlag.

Evrard, H. C. (2019). The Organization of the Primate Insular Cortex. Frontiers in Neuroanatomy, 13, 43. 10.3389/fnana.2019.00043

Fischl, B., van der Kouwe, A., Destrieux, C., Halgren, E., Ségonne, F., Salat, D. H., Busa, E., Seidman, L. J., Goldstein, J., Kennedy, D., Caviness, V., Makris, N., Rosen, B., & Dale, A. M. (2004). Automatically Parcellating the Human Cerebral Cortex. Cerebral Cortex, 14(1), 11–22. 10.1093/cercor/bhg087

Fabius, J. H., Moravkova, K., & Fracasso, A. (2022). Topographic organization of eye-position dependent gain fields in human visual cortex. Nature communications, 13(1), 7925.

Farrugia, C., Galdi, P., Irazu, I. A., Scerri, K., & Bajada, C. J. (2024). Local gradient analysis of human brain function using the Vogt-Bailey Index. Brain Structure and Function, 229(2), 497–512.

Fracasso, A., Dumoulin, S. O., & Petridou, N. (2021). Point-spread function of the BOLD response across columns and cortical depth in human extra-striate cortex. Progress in Neurobiology, 207, 102187. 10.1016/j.pneurobio.2021.102187

Fracasso, A., Luijten, P. R., Dumoulin, S. O., & Petridou, N. (2018). Laminar imaging of positive and negative BOLD in human visual cortex at 7 T. NeuroImage, 164, 100–111. 10.1016/j.neuroimage.2017.02.038

Fracasso, A., Van Veluw, S. J., Visser, F., Luijten, P. R., Spliet, W., Zwanenburg, J. J. M., Dumoulin, S. O., & Petridou, N. (2016a). Lines of Baillarger in vivo and ex vivo: Myelin contrast across lamina at 7 T MRI and histology. NeuroImage, 133, 163–175. 10.1016/j.neuroimage.2016.02.072

Fracasso, A., Van Veluw, S. J., Visser, F., Luijten, P. R., Spliet, W., Zwanenburg, J. J. M., Dumoulin, S. O., & Petridou, N. (2016b). Myelin contrast across lamina at 7T, ex-vivo and in-vivo dataset. Data in Brief, 8, 990–1003. 10.1016/j.dib.2016.06.058

Fukunaga, M., Li, T.-Q., van Gelderen, P., de Zwart, J. A., Shmueli, K., Yao, B., Lee, J., Maric, D., Aronova, M. A., Zhang, G., Leapman, R. D., Schenck, J. F., Merkle, H., & Duyn, J. H. (2010). Layer-specific variation of iron content in cerebral cortex as a source of MRI contrast. Proceedings of the National Academy of Sciences, 107(8), 3834–3839. 10.1073/pnas.0911177107

Gallay, D. S., Gallay, M. N., Jeanmonod, D., Rouiller, E. M., & Morel, A. (2012). The Insula of Reil Revisited: Multiarchitectonic Organization in Macaque Monkeys. Cerebral Cortex, 22(1), 175–190. 10.1093/cercor/bhr104

Gasquoine, P. G. (2014). Contributions of the insula to cognition and emotion. Neuropsychology review, 24(2), 77–87.

Gebhardt, S., & Nasrallah, H. A. (2023). The role of the insula in cognitive impairment of schizophrenia. Schizophrenia Research: Cognition, 32, 100277. 10.1016/j.scog.2022.100277

Geyer, S., Weiss, M., Reimann, K., Lohmann, G., & Turner, R. (2011). Microstructural parcellation of the human cerebral cortex–from Brodmann’s post-mortem map to in vivo mapping with high-field magnetic resonance imaging. Frontiers in human neuroscience, 5, 19.

Geyer, S., & Turner, R. (Eds.). (2013). Microstructural Parcellation of the Human Cerebral Cortex: From Brodmann’s Post-Mortem Map to in Vivo Mapping with High-Field Magnetic Resonance Imaging. Springer Berlin Heidelberg. 10.1007/978-3-642-37824-9

Glasser, M. F., & Essen, D. C. V. (2011). Mapping Human Cortical Areas In Vivo Based on Myelin Content as Revealed by T1- and T2-Weighted MRI. Journal of Neuroscience, 31(32), 11597–11616. 10.1523/JNEUROSCI.2180-11.2011

Glasser, M. F., Sotiropoulos, S. N., Wilson, J. A., Coalson, T. S., Fischl, B., Andersson, J. L., … & Wu-Minn HCP Consortium. (2013). The minimal preprocessing pipelines for the Human Connectome Project. Neuroimage, 80, 105–124.

Gogolla, N. (2017). The insular cortex. Current Biology, 27(12), R580–R586. 10.1016/j.cub.2017.05.010

Grady, L. J., & Polimeni, J. (2010). Discrete calculus: Applied analysis on graphs for computational science. Springer.

Han, X., Pham, D. L., Tosun, D., Rettmann, M. E., Xu, C., & Prince, J. L. (2004). CRUISE: cortical reconstruction using implicit surface evolution. NeuroImage, 23(3), 997–1012.

Hautus, M. J., Macmillan, N. A., & Creelman, C. D. (2021). Detection theory: A user’s guide. Routledge.

Inglese, M., Fleysher, L., Oesingmann, N., & Petracca, M. (2018). Clinical applications of ultra-high field magnetic resonance imaging in multiple sclerosis. Expert Review of Neurotherapeutics, 18(3), 221–230. 10.1080/14737175.2018.1433033

Isaacs, B. R., Mulder, M. J., Groot, J. M., Berendonk, N. van, Lute, N., Bazin, P.-L., Forstmann, B. U., & Alkemade, A. (2020). 3 versus 7 Tesla magnetic resonance imaging for parcellations of subcortical brain structures in clinical settings. PLOS ONE, 15(11), e0236208. 10.1371/journal.pone.0236208

Judas, M., & Cepanec, M. (2010). Oskar Vogt: The first myeloarchitectonic map of the human frontal cortex. Translational Neuroscience, 1, 72–94. 10.2478/v10134-010-0005-z

Klein, A., & Tourville, J. (2012). 101 Labeled Brain Images and a Consistent Human Cortical Labeling Protocol. Frontiers in Neuroscience, 6. 10.3389/fnins.2012.00171

Klugah-Brown, B., Wang, P., Jiang, Y., Becker, B., Hu, P., Uddin, L. Q., & Biswal, B. (2023). Structural–functional connectivity mapping of the insular cortex: A combined data-driven and meta-analytic topic mapping. Cerebral Cortex, 33(5), 1726–1738. 10.1093/cercor/bhac168

Koenig, S. H. (1991). Cholesterol of myelin is the determinant of gray-white contrast in MRI of brain. Magnetic Resonance in Medicine, 20(2), 285–291. 10.1002/mrm.1910200210

Kraff, O., Fischer, A., Nagel, A. M., Mönninghoff, C., & Ladd, M. E. (2015). MRI at 7 Tesla and above: Demonstrated and potential capabilities. Journal of Magnetic Resonance Imaging: JMRI, 41(1), 13–33. 10.1002/jmri.24573

Kurth, F., Eickhoff, S. B., Schleicher, A., Hoemke, L., Zilles, K., & Amunts, K. (2010a). Cytoarchitecture and Probabilistic Maps of the Human Posterior Insular Cortex. Cerebral Cortex, 20(6), 1448–1461. 10.1093/cercor/bhp208

Kurth, F., Zilles, K., Fox, P. T., Laird, A. R., & Eickhoff, S. B. (2010b). A link between the systems: Functional differentiation and integration within the human insula revealed by meta-analysis. Brain Structure and Function, 214(5–6), 519–534. 10.1007/s00429-010-0255-z

Lombaert, H., Grady, L., Polimeni, J. R., & Cheriet, F. (2011). Fast brain matching with spectral correspondence. Information Processing in Medical Imaging: Proceedings of the … Conference, 22, 660–673. 10.1007/978-3-642-22092-0_54

Lutti, A., Dick, F., fSereno, M., & Weiskopf, N. (2013). Using high-resolution quantitative mapping of R1 as an index of cortical myelination. NeuroImage, 93. 10.1016/j.neuroimage.2013.06.005

McGrath, C. L., Kelley, M. E., Holtzheimer III, P. E., Dunlop, B. W., Craighead, W. E., Franco, A. R., … & Mayberg, H. S. (2013). Toward a neuroimaging treatment selection biomarker for major depressive disorder. JAMA psychiatry, 70(8).

Medford, N., & Critchley, H. D. (2010). Conjoint activity of anterior insular and anterior cingulate cortex: Awareness and response. Brain Structure and Function, 214(5), 535–549. 10.1007/s00429-010-0265-x

Menon, V., Gallardo, G., Pinsk, M. A., Nguyen, V.-D., Li, J.-R., Cai, W., & Wassermann, D. (2020). Microstructural organization of human insula is linked to its macrofunctional circuitry and predicts cognitive control. eLife, 9, e53470. 10.7554/eLife.53470

Molnar-Szakacs, I., & Uddin, L. Q. (2022). Anterior insula as a gatekeeper of executive control. Neuroscience & Biobehavioral Reviews, 139, 104736. 10.1016/j.neubiorev.2022.104736

Nieuwenhuys, R. (2012a). The insular cortex. In Progress in Brain Research (Vol. 195, pp. 123–163). Elsevier. 10.1016/B978-0-444-53860-4.00007-6

Nieuwenhuys, R. (2012b). The Myeloarchitectonic Studies on the Human Cerebral Cortex of the Vogt-Vogt School, and Their Significance for the Interpretation of Functional Neuroimaging Data. Brain Structure & Function, 218. 10.1007/s00429-012-0460-z

Nomi, J. S., Schettini, E., Broce, I., Dick, A. S., & Uddin, L. Q. (2018). Structural Connections of Functionally Defined Human Insular Subdivisions. Cerebral Cortex, 28(10), 3445–3456. 10.1093/cercor/bhx211

Peerlings, J., Compter, I., Janssen, F., Wiggins, C. J., Postma, A. A., Mottaghy, F. M., Lambin, P., & Hoffmann, A. L. (2019). Characterizing geometrical accuracy in clinically optimised 7T and 3T magnetic resonance images for high-precision radiation treatment of brain tumours. Physics and Imaging in Radiation Oncology, 9, 35–42. 10.1016/j.phro.2018.12.001

Rolls, A. (2023). Immunoception: The insular cortex perspective. Cellular & Molecular Immunology. 10.1038/s41423-023-01051-8

Royer, J., Paquola, C., Larivière, S., Vos de Wael, R., Tavakol, S., Lowe, A. J., Benkarim, O., Evans, A. C., Bzdok, D., Smallwood, J., Frauscher, B., & Bernhardt, B. C. (2020). Myeloarchitecture gradients in the human insula: Histological underpinnings and association to intrinsic functional connectivity. NeuroImage, 216, 116859. 10.1016/j.neuroimage.2020.116859

Quabs, J., Caspers, S., Schoene, C., Mohlberg, H., Bludau, S., Dickscheid, T. and Amunts, K., 2022. Cytoarchitecture, probability maps and segregation of the human insula. Neuroimage, 260, p.119453

Saad, Z. S., & Reynolds, R. C. (2012). SUMA. NeuroImage, 62(2), 768–773. 10.1016/j.neuroimage.2011.09.016

Scholtens, L. H., de Reus, M. A., de Lange, S. C., Schmidt, R., & van den Heuvel, M. P. (2018). An MRI Von Economo – Koskinas atlas. NeuroImage, 170, 249–256. 10.1016/j.neuroimage.2016.12.069

Segerdahl, A. R., Mezue, M., Okell, T. W., Farrar, J. T., & Tracey, I. (2015). The dorsal posterior insula subserves a fundamental role in human pain. Nature neuroscience, 18(4), 499–500.

Stüber, C., Morawski, M., Schäfer, A., Labadie, C., Wähnert, M., Leuze, C., Streicher, M., Barapatre, N., Reimann, K., Geyer, S., Spemann, D., & Turner, R. (2014). Myelin and iron concentration in the human brain: A quantitative study of MRI contrast. NeuroImage, 93, 95–106. 10.1016/j.neuroimage.2014.02.026

Tanriover, N., Rhoton, A. L., Kawashima, M., Ulm, A. J., & Yasuda, A. (2004). Microsurgical anatomy of the insula and the sylvian fissure. Journal of Neurosurgery, 100(5), 891–922. 10.3171/jns.2004.100.5.0891

Tian, Y., & Zalesky, A. (2018). Characterizing the functional connectivity diversity of the insula cortex: Subregions, diversity curves and behavior. NeuroImage, 183, 716–733.

Tisserand, A., Philippi, N., Botzung, A., & Blanc, F. (2023). Me, Myself and My Insula: An Oasis in the Forefront of Self-Consciousness. Biology, 12(4), 599. 10.3390/biology12040599

Trattnig, S., Springer, E., Bogner, W., Hangel, G., Strasser, B., Dymerska, B., Cardoso, P. L., & Robinson, S. D. (2018). Key clinical benefits of neuroimaging at 7T. NeuroImage, 168, 477–489. 10.1016/j.neuroimage.2016.11.031

Türe, U., Yaşargil, M. G., Al-Mefty, O., & Yaşargil, D. C. H. (2000). Arteries of the insula. Journal of Neurosurgery, 92(4), 676–687. 10.3171/jns.2000.92.4.0676

Uddin, L. Q., Kinnison, J., Pessoa, L., & Anderson, M. L. (2014). Beyond the Tripartite Cognition–Emotion–Interoception Model of the Human Insular Cortex. Journal of Cognitive Neuroscience, 26(1), 16–27. 10.1162/jocn_a_00462

Uddin, L. Q., Nomi, J. S., Hébert-Seropian, B., Ghaziri, J., & Boucher, O. (2017). Structure and Function of the Human Insula. Journal of Clinical Neurophysiology: Official Publication of the American Electroencephalographic Society, 34(4), 300–306. 10.1097/WNP.0000000000000377

van Dijk, J. A., Fracasso, A., Petridou, N., & Dumoulin, S. O. (2021a). Laminar processing of numerosity supports a canonical cortical microcircuit in human parietal cortex. Current Biology, 31(20), 4635–4640.

van Dijk, J. A., Fracasso, A., Petridou, N., & Dumoulin, S. O. (2021b). Validating linear systems analysis for laminar fMRI: Temporal additivity for stimulus duration manipulations. Brain Topography, 34(1), 88–101.

Vogt, O. (1903). Zur anatomischen Gliederung des Cortex cerebri. J Psychol Neurol, 2(4), 160–180.

Waehnert, M. D., Dinse, J., Schäfer, A., Geyer, S., Bazin, P.-L., Turner, R., & Tardif, C. L. (2016). A subject-specific framework for in vivo myeloarchitectonic analysis using high resolution quantitative MRI. NeuroImage, 125, 94–107. 10.1016/j.neuroimage.2015.10.001

Waehnert, M. D., Dinse, J., Weiss, M., Streicher, M. N., Waehnert, P., Geyer, S., Turner, R., & Bazin, P.-L. (2014). Anatomically motivated modeling of cortical laminae. NeuroImage, 93, 210–220. 10.1016/j.neuroimage.2013.03.078

