## Supplementary Material for "Delineating In-Vivo T1-Weighted Intensity Profiles Within the Human Insula Cortex Using 7-Tesla MRI"

**AUTHOR BLOCK: *C. DALBY**[1], A. DIBBLE[1], J. CARVALHEIRO[1], F. QUEIRAZZA[2], M. SEVEGNANI[3], M. HARVEY[1], M. SVANERA[1], A. FRACASSO[1]
[1]School of Psychology & Neuroscience, Univ. of Glasgow, Glasgow, United Kingdom;
[2]School of Health & Wellbeing, Univ. of Glasgow, Glasgow, United Kingdom;
[3]School of Computing Science, Univ. of Glasgow, Glasgow, United Kingdom

**Corresponding author**
Alessio Fracasso
Hillhead Street 62
G12 8QE
University of Glasgow, Scotland UK


### Removing Cortical Curvature and Cortical Thickness Contributions

The human cortex is characterised by a complex pattern of concavity and convexity, and myelin content varies significantly with local cortical convexity (Annese et al. 2004). T1-w signal is known to vary systematically with cortical curvature and thickness, convex regions tend to be more myelinated than concave regions and vice-versa (Sereno et al., 2013). Removing the contribution of local curvature and cortical thickness was necessary because both have been shown to influence T1-w intensity changes in in-vivo imaging (Dumoulin et al., 2018; Fracasso, Petridou & Dumoulin, 2016; Lutti et al., 2013; Sereno et al., 2013), as well as local myelination in ex-vivo studies (Annese et al., 2004). It is common practice to remove curvature and thickness contributions from the T1-w signal prior to further analysis (Sereno et al., 2013), as the relationship between T1-w intensity and local curvature may obscure global changes in T1-w intensity.
Without this step, the clustering analysis would likely capture known curvature-related differences in T1-w intensity between the bottom of the sulci and the top of the gyri within the insula, thereby masking other relevant T1-w intensity variations across the ROI (Sereno et al., 2013; Lutti et al., 2013). We removed the contributions of local cortical curvature and thickness for each level-set along cortical depth (8 level-sets overall from white matter to grey matter surface). We fit a linear model for each level-set where we factored the contribution of local cortical curvature and local cortical thickness plus the interaction between the two. For each level-set along cortical depth, we used the residuals of the linear model just described, after adding the estimated intercept T1-w signal.
All analyses were performed at an individual participant, individual insula (left/right) level, thus addressing variations in insular folding across participants.

**Individual-Level T1-w Intensity Profiles and Clusters from the Desikan-Killiany Atlas (DKT) and Von Economo-Koskinas (VKA) Insula ROI, AHEAD Dataset**

Our analysis revealed distinct high- and low-T1-w intensity profiles across DTK insula ROI at the individual-level (Figure SF1). T1-w signal monotonically decreases along cortical depth in both the high- and low-T1-w signal cluster. We evidence two distinct high- and low-T1-w intensity profiles when employing the DKT and VKA atlas for the same individual (Figure SF1A&B). We show the same results in a second participant to exemplify the robustness of the findings (Figure SF1C, D).

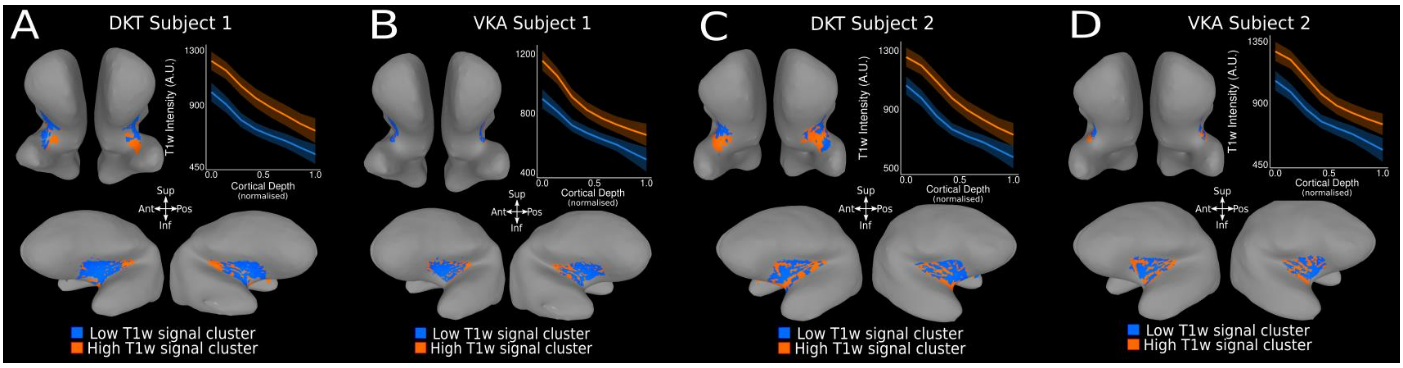

***SF1: Supplementary Figure SF1, T1-w Intensity Profiles and Clusters in DTK and VKA insula ROI for AHEAD Dataset Individuals.***

***A.** Average high (orange) and low (blue) T1-w intensity profiles for the insula DKT ROI at the individual subject level. T1-w intensity is plotted along cortical depth and clusters are visualised on the MNI SUMA surface of the individual's cortex (high T1-w intensity cluster – orange – and low T1-w intensity cluster – blue – ). **B.** Average high and low T1-w intensity profiles for the insula VKA ROI in the same subject. **C-D.** Replication of A and B in a second subject example.*

**DKT Atlas, The Influence of The Middle Cerebral Artery on T1-w Signal Intensity**

The middle cerebral artery (MCA) is a major artery supplying blood to the lateral portion of the cerebral cortex, including the primary motor and sensory areas. Originating from the internal carotid artery, it runs laterally through the Sylvian fissure, closely adjacent to the insula cortex (Medrano-Martorell et al., 2021). The M2 segment of the middle cerebral artery refers to the portions of this artery that course laterally beyond the insula, branching extensively within the Sylvian fissure. This segment represents the continuation of the MCA after it traverses the surface of the insula, where it bifurcates or trifurcates into smaller arteries that supply cortical regions adjacent to and beyond the insula (Tanriover et al., 2004). Thus, while the M2 segment directly follows the insular traversal of the MCA, it is primarily involved with vascularising areas external to the insula itself (Türe et al., 2000).

For the DKT atlas, we assessed the influence of MCA proximity to our clustering solution using a linear model, fitting the relationship between T1-w signal and normalised cortical depth, using cluster location as a categorical variable. We test the differences between separate clusters in terms of intercept (T1-w value at the white matter interface) and slope (rate of change of T1-w signal rate per unit change of cortical depth).

Crucially, large arteries as MCA and their associated branches appear as bright tubular structures in T1-w images. Such a high T1-w signal intensity in proximity with the superficial grey matter border could bias T1-w profiles. Segmenting out the MCA could be a viable approach to mitigate this bias; however, due to potential partial volume effects in the T1-w image, segmenting out the MCA would not fully eliminate the MCA's high T1-w intensity influencing neighbouring voxels within adjacent grey matter. This influence could potentially affect the estimation of T1-w profiles across cortical depth.
For this reason, rather than segmenting out the MCA, we explicitly analysed T1-w profiles close to the MCA to directly assess the potential influence of partial volume effects on our measurements. To assess the influence of MCA in the T1-w signal profiles we display the grey matter location of MCA with respect to our clustering results and statistically assess differences in our clustering results that might be ascribed to MCA proximity.

The profiles in the high T1-w signal cluster can be found in two separate cortical locations along the DKA insula ROI: in the posterior-superior portion and the anterior-inferior portion of the DKA insula ROI (Figure SF2). Importantly, the anterior-inferior section of the DKA insula ROI is situated near the MCA artery. Here, we report the grey matter location of the high- and low- T1-w signal cluster in individual subjects, as well as the average T1-w profile for the posterior-superior, middle and anterior-inferior T1-w signal insular clusters (Figure SF2A-D).

We quantitatively compared the high-T1-w signal cluster in the posterior-superior and anterior-inferior DKA insula ROI. Statistical analysis shows a difference in intercept (estimate = 87.80, $t=12.64$, $p<0.001$) and slope (estimate =-195.16 $t=-16.79$, $p<0.001$), indicating a shallower slope for the anterior-inferior high-T1-w signal cluster compared to the posterior-superior (Figure SF2D). This is to be expected as partial volume influence from the MCA can contribute to increasing T1-w signal close to the pial surface for the anterior-inferior T1-w signal cluster. However, although the average T1-w profile in the anterior portion of the DKT insula ROI is statistically different from its posterior counterpart, visual inspection of the profiles (Figure SF2) indicates that the presence of MCA does not disrupt the overall shape of T1-w signal profiles along cortical depth in the anterior high-T1-w signal cluster, and the high-T1-w cluster in the anterior insula is not driven by the MCA proximity.

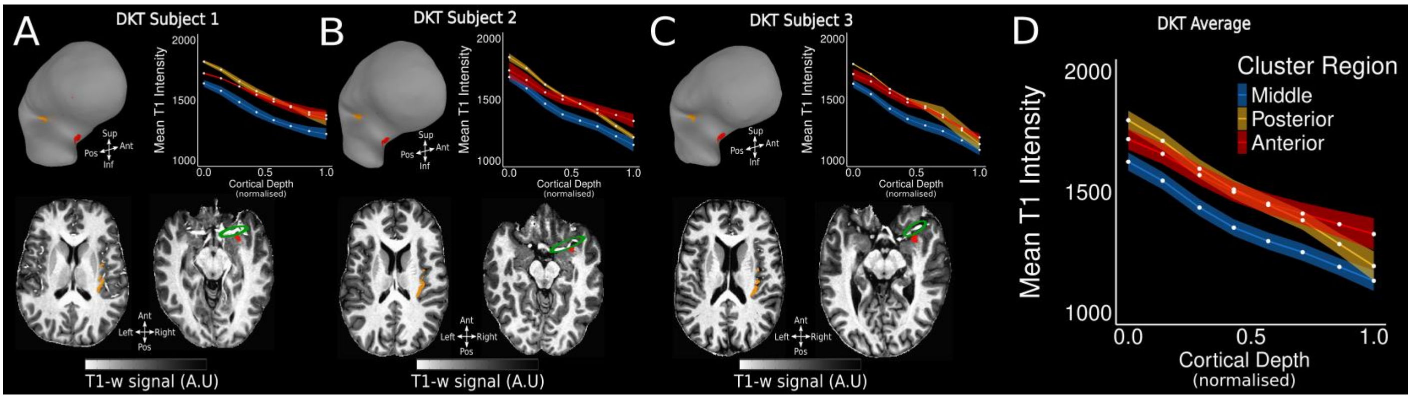

***SF2: Supplementary Figure 2, Proximity of the Anterior-Inferior High T1-w Cluster to Middle Cerebral Artery.***

***A-C.*** *T1-w signal clusters, located in the posterior-superior (yellow), anterior-inferior (red) and middle (blue) region of the DKT insula ROI across 3 subjects. High T1-w signal clusters are visualised on the right hemisphere SUMA MNI surface (panel 1) and T1-w intensity profiles are plotted along cortical depth (panel 2). High T1-w signal clusters are visualised on each subject's T1-w volume (panel 3-4). Green circle indicates the location of the middle cerebral artery segment.* ***D.*** *Average T1-w intensity profiles for the posterior-superior (yellow), anterior-inferior (red) and middle (blue) region of the DKT insula ROI for 21 subject (Glasgow Dataset) plotted across cortical depth.*

**Clusters or Gradients, simulations**

In any clustering problem, it is important to ask how to characterize neighbouring locations that differ with respect to a given metric. In our case, it is possible to hypothesize that neighbouring cortical locations could be represented by two separate clusters (Figure SF3A) or, alternatively by a gradient (Figure SF3B), smoothly transitioning between the neighbouring cortical locations.

'True' separate clusters would be separated by a sudden, transient change in properties when moving from one cortical location to another. In contrast, a gradual change implies a smooth transition without abrupt shifts in the measured properties.

Crucially, while a clustering solution - corroborated by silhouette scores - may partition the parameter space between two neighbouring regions, it does not inherently reveal whether the underlying properties are organized as genuinely distinct clusters (with transient changes – a boundary between the clusters) or as a continuous gradient. This distinction can be illustrated using simulated examples: one scenario with two discrete clusters along an arbitrary dimension (Figure SF3A), and another with a continuous gradient along the same dimension (Figure SF3B).

The simulation shows how a kmeans solution per se is not informative regarding the trends present in the underlying distributions. Separate clusters can still be identified in the presence of a continuous gradient. It is possible to disentangle between 'separate clusters' and the 'smooth gradient' hypothesis by comparing the fits between alternative functions along the dimension of interest (in our manuscript: cortical distance).

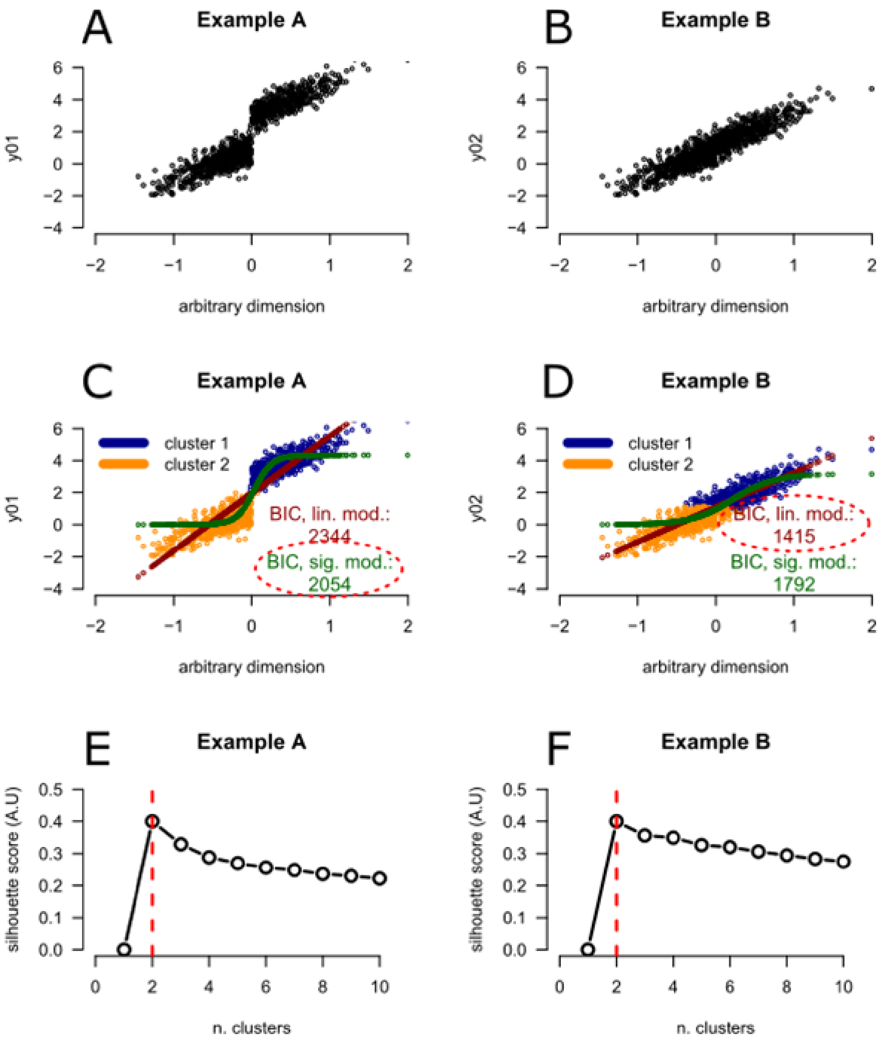

*SF3: **Supplementary Figure 3.** here we show two simulated examples. **Panel A:** Example A with two discrete clusters along an arbitrary dimension x and **Panel B:** Example B with a continuous gradient along the same dimension. In **Panel C** we report the kmeans solution of Example A with 2 clusters (cluster 1 and cluster 2), highlighting a separation between the two clouds of points in the parameter space. We also report the results of two models fit to the data: the best fitting linear fit (red points) and the best fitting cumulative Gaussian fit (green points), along with the BIC criterion for the two models. Please note how the cumulative Gaussian fit yields a better fit (lower BIC value, red dashed circle), compared to the linear fit, indicating that a transient change better captures the trend of the y01 variable along the arbitrary dimension x. **Panel D** shows similar information as in Panel C, but this time for Example B, in the presence of a smooth transition of the variable y02 along the arbitrary dimension x. Please note how in this case the cumulative Gaussian fit yields a worse fit (higher BIC value), compared to the linear fit (red dashed circle), indicating that a gradual change better capture the trend of the y02 variable along the arbitrary dimension x, rather than a transient change. **Panels E-F.** Silhouette scores from Example A and B, respectively. Crucially, in both cases (Example A and B) a cluster solution with 2 clusters is identified as the optimal (red dashed line). Although the silhouette score profile is clearly steeper in Example A (separate clusters) compared to Example B (smooth gradient), as could be expected. Overall, this simulation shows how a kmeans solution, corroborated by silhouette scores, is not informative per se regarding the trends present in the underlying*

*distributions. Separate clusters can still be identified in the presence of a continuous gradient. The discriminant between 'separate clusters' and 'smooth gradient' can be achieved by comparing the fits between alternative functions along the dimension of interest (in our manuscript: cortical distance).*

The analysis presented in SF3 and the analysis presented in Figure 6 of the main manuscript shows how the clusters identified by the clustering analysis are indeed indicative of a transient change in properties along cortical distance in the human insula, not a simple gradient along cortical space.

**Functional gradients in the human insula**

Tian & Zalesky (2018) and Farrugia and colleagues (2024), proposed methods to identify and quantify meaningful clusters / gradients along the cortical surface. Interestingly Tian & Zalesky (2018) applied these methods to the study of the human insula and identified meaningful gradients along the insula anterior posterior dimension based on the analysis of human resting state data (functional MRI) and dimensionality reduction.
Tian and Zalesky (2018) propose an approach based on a PCA dimensionality reduction and using the first 5 principal components to compute the connectivity between all insula voxels. Subsequently the authors successfully identified the anterior-posterior direction as the main direction along which features change along the human insula.
Farrugia and colleagues used the Vogt-Bailey index (VB), which represents the normalised connectivity of the graph Laplacian of given input features, used to describe the extent of feature similarity. Of particular interest within this context is the searchlight VB Index (Bajada et al., 2020), which computes a VB Index per vertex based on the neighbourhood data of directly adjacent vertices and can highlight effective 'boundaries' between otherwise homogeneous cortical locations.
Both manuscripts (Farrugia et al., 2024 and Tian & Zalesky, 2018) apply their method to human resting state data. This type of data shows a high degree of variability and a relatively low degree of collinearity, that is: the correlation between time series of different voxels is relatively low (in the absence of highly correlated physiological noise). This is an important detail as one fundamental step of both methods is computing the Laplacian matrix and its eigenvectors, characterizing the connectivity between voxels in the dataset. The Laplacian consists in the difference between a *degree matrix* and an *adjacency matrix* (Bajada et al., 2020).

*Adjacency matrix:* The adjacency matrix is a square matrix (i.e., the same number of rows and columns) where every row and every column represent a single node (voxel), and the elements in the matrix represent the relationships between the row node and the column node. For example, either dot product or the correlation between voxels.

*Degree matrix:* the entries along the diagonal represent the degree of each node, that is, the number of nodes that are connected (adjacent) to that node.

A stable computation of the eigenvector depends on the collinearity of the input data, particularly in the adjacency matrix. High collinearity can introduce problems related to numerical instability: the matrix become nearly singular and thus very sensitive to small

perturbations. This can lead to inaccurate or unstable eigenvalue computations due to rounding errors.
Structural cortical depth dependent data, as the one presented in our manuscript, is highly collinear in nature (the correlation between cortical depth dependent profiles is relatively high), due to the highly stereotypical increasing or decreasing (depending on the MRI contrast) trends of structural MRI signal along cortical depth. Thus, it not straightforward to predict how the methods proposed by Farrugia et al. (2024) and Tian & Zalesky (2018) would operate on data as the one presented in our manuscript. However, it is important to acknowledge that these methods could in principle be applied and provide useful insights regarding the problem of parcellating the human insula, in-vivo.
Kmeans can also be affected by highly collinear input, however the algorithm is sensitive to absolute values, and can capture offset differences between profiles, as those presented in the main manuscript.

**References**

Annese, J., Pitiot, A., Dinov, I. D., & Toga, A. W. (2004). A myelo-architectonic method for the structural classification of cortical areas. Neuroimage, 21(1), 15-26.

Bajada, C. J., Campos, L. Q. C., Caspers, S., Muscat, R., Parker, G. J., Ralph, M. A. L., ... & Trujillo-Barreto, N. J. (2020). A tutorial and tool for exploring feature similarity gradients with MRI data. *NeuroImage*, *221*, 117140.

Dumoulin, S. O., Fracasso, A., van der Zwaag, W., Siero, J. C. W., & Petridou, N. (2018). Ultra-high field MRI: Advancing systems neuroscience towards mesoscopic human brain function. NeuroImage, 168, 345–357. https://doi.org/10.1016/j.neuroimage.2017.01.028

Farrugia, C., Galdi, P., Irazu, I. A., Scerri, K., & Bajada, C. J. (2024). Local gradient analysis of human brain function using the Vogt-Bailey Index. *Brain Structure and Function*, *229*(2), 497-512.

Fracasso, A., Petridou, N., & Dumoulin, S. O. (2016). Systematic variation of population receptive field properties across cortical depth in human visual cortex. Neuroimage, 139, 427-438.

Lutti, A., Dick, F., fSereno, M., & Weiskopf, N. (2013). Using high-resolution quantitative mapping of R1 as an index of cortical myelination. NeuroImage, 93. https://doi.org/10.1016/j.neuroimage.2013.06.005

Medrano-Martorell, S., Pumar-Pérez, M., González-Ortiz, S., & Capellades-Font, J. (2021). A review of the anatomy of the middle cerebral artery for the era of thrombectomy: A radiologic tool based on CT angiography and perfusion CT. Radiología (English Edition), 63(6), 505–511. https://doi.org/10.1016/j.rxeng.2021.10.004

Tanriover, N., Rhoton, A. L., Kawashima, M., Ulm, A. J., & Yasuda, A. (2004). Microsurgical anatomy of the insula and the sylvian fissure. Journal of Neurosurgery, 100(5), 891–922. https://doi.org/10.3171/jns.2004.100.5.0891

Tian, Y., & Zalesky, A. (2018). Characterizing the functional connectivity diversity of the insula cortex: Subregions, diversity curves and behavior. NeuroImage, 183, 716-733.

Türe, U., Yaşargil, M. G., Al-Mefty, O., & Yaşargil, D. C. H. (2000). Arteries of the insula. Journal of Neurosurgery, 92(4), 676–687. https://doi.org/10.3171/jns.2000.92.4.0676

Sereno, M. I., Lutti, A., Weiskopf, N., & Dick, F. (2013). Mapping the human cortical surface by combining quantitative T 1 with retinotopy. Cerebral cortex, 23(9), 2261-2268.
